# Hierachical Resampling for Bagging in Multi-Study Prediction with Applications to Human Neurochemical Sensing

**DOI:** 10.1101/856385

**Authors:** Gabriel Loewinger, Prasad Patil, Kenneth T. Kishida, Giovanni Parmigiani

## Abstract

We propose the “study strap ensemble,” which combines advantages of two common approaches to fitting prediction models when multiple training datasets (“studies”) are available: pooling studies and fitting one model versus averaging predictions from multiple models each fit to individual studies. The study strap ensemble fits models to bootstrapped datasets, or “pseudo-studies.” These are generated by resampling from multiple studies with a hierarchical resampling scheme that generalizes the randomized cluster bootstrap. The study strap is controlled by a tuning parameter that determines the proportion of observations to draw from each study. When the parameter is set to its lowest value, each pseudo-study is resampled from only a single study. When it is high, the study strap ignores the multi-study structure and generates pseudo-studies by merging the datasets and drawing observations like a standard bootstrap. We empirically show the optimal tuning value often lies in between, and prove that special cases of the study strap draw the merged dataset and the set of original studies as pseudo-studies. We extend the study strap approach with an ensemble weighting scheme that utilizes information in the distribution of the covariates of the test dataset.

Our work is motivated by neuroscience experiments using real-time neurochemical sensing during awake behavior in humans. Current techniques to perform this kind of research require measurements from an electrode placed in the brain during awake neurosurgery and rely on prediction models to estimate neurotransmitter concentrations from the electrical measurements recorded by the electrode. These models are trained by combining multiple datasets that are collected *in vitro* under heterogeneous conditions in order to promote accuracy of the models when applied to data collected in the brain. A prevailing challenge is deciding how to combine studies or ensemble models trained on different studies to enhance model generalizability.

Our methods produce marked improvements in simulations and in this application. All methods are available in the `studyStrap` CRAN package.

## 1. Introduction

### 1.1. Multi-Study Learning

It has become increasingly common to encounter settings where multiple datasets (or “studies”) with common covariates and outcomes are available for training prediction models. These opportunities arise, for example, in genetics (Zhang et al., 2020), neuroimaging (Glocker et al., 2019) and HIV (Ramon, Belanche-Muñoz and Pérez-Enciso, 2019). Leveraging training information from multiple datasets is desirable since algorithms trained on data from one study often perform poorly when used for the same prediction task in other studies (Patil and Parmigiani, 2018). Developing algorithms that explicitly account for the heterogeneity across datasets is critical to generating prediction models that are replicable across settings.

Poor out-of-sample generalizability can result from different sources of “dataset shift,” a discrepancy in the distribution of training and test data (Kouw and Loog, 2019; Yang et al., 2020). Dataset shift can take different forms: 1) “virtual drift” or “covariate shift,” occurs when there are differences in the distribution *f* (𝕏) of the covariates; 2) “concept shift” arises when the conditional probability *f* (**y** | (𝕏) changes; 3) “hybrid shift” presents when both *f* (**y** | 𝕏) and *f* (𝕏) change.

Methods proposed in the transfer learning literature aim to improve performance of supervised learning in a new dataset (“domain”). Multi-source domain adaptation methods leverage multiple domains to improve performance on a target dataset using available information on its covariates (Kouw and Loog, 2019), while domain generalization uses data from multiple domains to make a model more generalizable to an unseen dataset (Wang et al., 2021). In the present work, we focus on domain generalization and multi-source domain adaptation problems (Sun, Shi and Wu, 2015) motivated by neuroscience applications. Extensive literature on transfer learning and domain adaptation work has proposed approaches to reweighting samples (Shimodaira, 2000; Sugiyama et al., 2008) to align the distribution of data (e.g., the marginal the distribution of the covariates) (Kouw and Loog, 2019; Sun, Shi and Wu, 2015). Here we propose hierarchical resampling techniques coupled with covariate distribution-based weighting schemes towards similar ends. We leverage ideas from the rich literature in transfer learning and the rapidly growing “multi-study” statistics perspective that proposes methods to combine studies in supervised (Guan, Parmigiani and Patil, 2020; Ramchandran, Patil and Parmigiani, 2020; Ren et al., 2021), unsupervised (De Vito et al., 2019; Roy et al., 2019) and inference settings (Guo et al., 2021; Rashid et al., 2020).

Multi-study learning methods seek to leverage information from multiple studies to improve the replicability of models. A standard approach in multi-study settings is to simply pool studies together and fit a single model on this merged dataset (Guan, Parmigiani and Patil, 2020; Sun, Shi and Wu, 2015), which we refer to as “Training On the Merged Dataset” algorithm, or “TOM”. While the larger sample size and the simplicity of the approach are attractive, this procedure can result in poor out-of-sample prediction performance when there is high between-study heterogeneity in the joint distribution of the covariates or in the conditional distribution of the outcome given the covariates. Ensembling, or combining predictions from models trained on different studies rather than combining the study data themselves, has been shown to be a useful framework for accounting for heterogeneity and simultaneously borrowing strength across different datasets (Guan, Parmigiani and Patil, 2020; Guo, Shah and Barzilay, 2018; Patil and Parmigiani, 2018; Ramchandran, Patil and Parmigiani, 2020; Sun, Shi and Wu, 2015). A simple form is the “Observed-Studies Ensemble” (“OSE”), in which a model is fit on each study separately and then the predictions from all the models are aggregated through a weighted average.

### 1.2. A Multi-study Challenge in Neurochemical Sensing

Studying the neurobiological underpinnings of decision making is critical for developing treatments for neurological and psychiatric conditions such as drug addiction and Alzheimer’s disease. To study the brain circuitry involved in diseases of the nervous system, neuroscientists often seek to measure changes in levels of neurotransmitters (e.g., dopamine) used to relay messages between brain cells. Historically such experiments have mostly been conducted in non-human model organisms (Volkow, Wise and Baler, 2017). Until recently, however, brain researchers have not had the tools necessary to measure fluctuations in neurotransmitter levels in humans on a time scale rapid enough to enable investigating the link between these brain signals and real-time changes in behavior and cognition (Kishida et al., 2016). Fast Scan Cyclic Voltammetry (FSCV) is an invasive electrochemical technique that allows for the estimation of neurotransmitter changes at a rapid time scale (10 measurements per second) in awake humans who are performing decision-making tasks. Historically used in rodent models, FSCV technology for humans has recently been developed (Bang et al., 2020; Kishida et al., 2016; Moran et al., 2018), offering unprecedented opportunity to monitor human neurochemical levels that is currently impossible to estimate with other common human neuroscience techniques such as non-invasive brain imaging (e.g., Functional Magnetic Resonance Imaging (fMRI) or Positron Emission Tomography (PET)).

FSCV presents, however, statistical challenges that must be resolved to ensure the generation of accurate neurotransmitter estimates. This is because this approach inherently relies on statistical models to translate raw measurements into estimates of neurotransmitter concentration. Briefly, the technique functions by varying the voltage potential on an electrode and measuring the resultant changes in electrical current. The recordings produce a high dimensional time series signal. The vector of current measurements at each time point can be used as covariates to predict neurotransmitter concentration. As FSCV does not directly measure this outcome, one must train models on datasets generated in vitro, where a ground truth (i.e., the true concentrations) is known.

Models trained in this manner often suffer, however, from poor generalizability, as the environments in which in vitro datasets are constructed differ markedly from that of the brain. To improve the generalizability of the models, it is common to train them on a dataset that pools together multiple in vitro datasets (referred to as “calibration” datasets in the FSCV literature), each generated on a different electrode (Bang et al., 2020; Kishida et al., 2016; Moran et al., 2018). In practice, this is a multi-study learning problem. While different in vitro datasets share the same outcome and covariates, electrodes are hand-made and differ subtly both in their general physical properties (e.g., the length of the electrode) and in the properties of their recorded signals (i.e., their electrical responses). As a result, data collected using different electrodes exhibit considerable heterogeneity both in the marginal distribution of the covariates (i.e., the magnitude of current measurements recorded at a fixed set of voltage potentials) and in the conditional distributions of the outcome (neurotransmitter concentration) given the covariates. Treating all observations made on one electrode as a study allows for a systematic approach to account for the fact that no one in vitro dataset will exactly capture the properties of data collected in the brain. Indeed the original work to implement FSCV in humans (Kishida et al., 2016) reports that training models on datasets that combine “studies” (i.e., in vitro datasets) of multiple electrodes substantially improves the cross-electrode generalizability and accuracy of estimates.

### 1.3. A General Framework for Multi-Study Training

In our neurochemical sensing application, as well as other contexts (Ventz, Mazumder and Trippa, 2020), OSE and TOM each outperform each other at times, but it can be difficult to predict under which conditions one will be superior. We sought to create an encompassing framework combining advantages of both approaches: 1) training models on datasets that combine observations from multiple studies and 2) ensembling to improve out-of-sample performance.

We propose a method that generates an artificial collection of “studies” by resampling the set of the observed studies in a manner that is useful for multi-study ensemble learning. We term such collection a “study strap replicate” and each member a “pseudo-study.” We refer to the original studies, without any resampling, as “observed studies” and the resampling procedure as the “study strap.” Each pseudo-study can then be used as a training dataset to fit a prediction model. In a study strap replicate, each pseudo-study includes observations from a subset of the observed studies, in different proportions. While one could generate pseudo-studies by resampling from the merged dataset with replacement (i.e., standard “bagging” or “bootstrap aggregation” (Breiman, 1996a)), this would produce many bootstrap datasets that have observations from all or many of the observed studies. Conversely, resampling from each of the observed studies separately would result in pseudo-studies that only have observations from a single study. To control the between pseudo-study heterogeneity in our resampling approach, we randomly determine the number of observations to resample from each observed study via a multinomial draw. For example, say we have 5 observed studies, each with 100 observations. A pseudo-study could be constructed by drawing 30 observations from study 1, 10 from study 3, and 60 from study 5. We will analytically demonstrate that special cases of the study strap generate both the set of observed studies and the merged dataset as as “pseudo-studies.”

Our proposed study strap approach leverages the concept of bagging, which implements bootstrap resampling in prediction settings (Breiman, 1996a). Training one or more learners on each of several bootstrap samples and then ensembling the resulting models can enhance performance, often through a reduction in variance. We also build on an existing hierarchical resampling approach (Davison and Hinkley, 1997) (pp.100-102): the *randomized cluster bootstrap*, where both clusters and observations within a cluster are resampled *with* replacement. As we discuss below, standard (non-hierarchical) bagging of the merged dataset as well as the randomized cluster bootstrap are special cases of the study strap resampling scheme.

In our neurochemical sensing application, models are trained after the covariates of the test set (signals recorded in the brain) have been observed. Past FSCV work has reported that using training sets where the joint distribution of the covariates is similar to that of the test set improves prediction performance (Kishida et al., 2016). Motivated by this observation, we propose to upweight models trained on datasets that have similar covariate profiles to that of the target task or population. We extend this concept to the study strap by only training models to be included as members of an ensemble if the corresponding pseudo-studies share a covariate distribution similar to that of the test set.

## 2. Methods

### 2.1. Notation and Problem Statement

We consider *K* training (observed) studies with common outcomes and covariates, and make predictions on data collected in a separate study (study *K* + 1). The observed training studies are noted by {𝕊_1_, …, 𝕊_*K*_} where 𝕊_*k*_ = [**y**_*k*_ | 𝕏_*k*_], **y**_*k*_ is the outcome variable in the *k*^*th*^ study and 𝕏_*k*_ is its design matrix. The studies have sample sizes *n*_1_, …, *n*_*K*_ and we define *N* = Σ_*k*_ *n*_*k*_. We denote the set {1, 2, …, *K*} as [*K*]. We assume the data from each study are generated independently *across* studies. The covariates from study *k* are drawn from the distribution 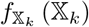, and the outcome from 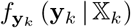. Subscripts will be dropped when we can do so without ambiguity. We allow for between-study heterogeneity, allowing both the underlying marginal distributions, *f* (𝕏_*k*_), *f* (**y**_*k*_) and the conditional distribution, *f* (**y**_*k*_ | 𝕏_*k*_) to vary across studies. These are known in the dataset shift literature as “covariate shift” and “concept drift” respectively (Yang et al., 2020).

We aim to make predictions for target study *K* + 1, possibly using its design matrix, 𝕏_*K*+1_. We propose methods for 1) *general* prediction on an exchangeable study without knowledge of the covariate profile and 2) *customized* prediction on an exchangeable study, with knowledge of the covariate profiles of all samples (or a representative subsample) in the new (target) study. In 1) {𝕊_1_, …, 𝕊_*K*_} are observed. In 2) {𝕏_*K*+1_, 𝕊_1_, …, 𝕊_*K*_} are observed.

Denote a model trained on data from the *k*^*th*^ study by **Ŷ**_*k*_(·) and the predictions made with this model on the covariates of study *K* + 1 by **Ŷ**_*k*_ (𝕏_*k*+1_). When we discuss ensembles, we express them in terms of the original models. For example, the observed-studies ensemble **Ŷ**_*OSE*_ (·) trains a single model on each study and then ensembles the resulting models:

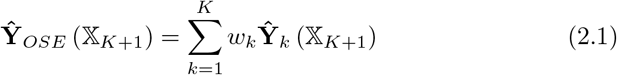

where *w*_*k*_ is the weight on the predictions of the *k*^*th*^ model.

### 2.2. Multi-study Ensembling via Stacking

One approach for determining the weights in (2.1) is the multi-study counterpart of stacking (Breiman, 1996b) introduced in Patil and Parmigiani (2018). First a model is fit on each observed study. Then a regression is fit to **y** and 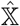 defined as

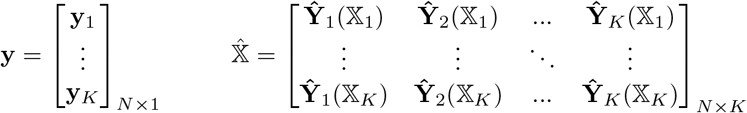

using non-negative least squares (NNLS). The weights *ŵ* are the NNLS coefficient estimates. In multi-study prediction, this type of stacking rewards cross-study prediction performance. We use it in several cases, and include implementation details in Supplementary Section A.3.

### 2.3. Study Strap

We propose the *study strap*, a general approach to generating collections of *pseudo-studies* for both general and customized prediction. The pseudo-studies serve as training datasets to which any statistical learning approach can be applied. The study strap is a hierarchical resampling scheme which generates each pseudo-study by first selecting the proportion of observations to resample from each of the observed studies (and implicitly which observed studies to resample from) in a “study-level” resampling step. Then in the “observation-level” step, individual observations are resampled from each observed study (with or without replacement) according to the proportions drawn in the first step. The first stage of the resampling procedure is controlled by *b*, the “bag size” tuning parameter. The sample size of a psuedo-study depends on the sample sizes of the observed studies, *n*_1_, …, *n*_*K*_.

Formally, in pseudo-code, for pseudo-study *r*, `round` 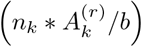 observations are resampled from observed study *k*, where 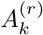 is the *k*^*th*^ element of the multinomial draw 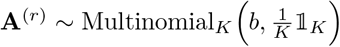. The function `round`(·) maps a real number to the closest integer. We refer to **A**^(*r*)^ as the “study bag.” *b*, the “bag size” tuning parameter, is used to control the degree to which multiple studies contribute observations to the composition of a pseudo-study. When *b* = 1, all observations for a given pseudo-study are drawn from one observed study. When *b* ≤ *K, b* is the maximum number of observed studies that could be selected to contribute observations to a pseudo-study. When *b* = *K*, the study strap draws pseudo-studies similarly to the randomized cluster bootstrap. As *b* grows large, each of the observed studies will tend to be represented (see Supplement Proposition 7) and contribute equal proportions of their sample size to a given pseudo-study: 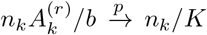 (for fixed *n*_*k*_) by the Weak Law of Large Numbers. The number of observed studies that contribute to a given pseudo-study, 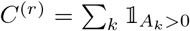, is random with a distribution that depends on *b*. We show in the Supplement (Proposition 6) that this distribution has the form: 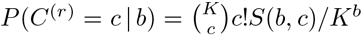, where *S*(*b, c*) is Sterling’s number of the second kind.

The bag size also determines the proportion of observations that can be resampled from a study. Since we draw `round` 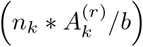 observations from the *k*^*th*^ study, 1*/b* is interpretable as the smallest non-zero proportion of observations one can resample from any of the *K* studies (i.e., `round`(*n*_*k*_*/b*) is the smallest number of (non-zero) observations that could be be drawn from study *k*). Described another way, at each iteration, the *study strap* randomly cuts each study into *b* roughly equally sized pieces. It then draws an **A**^(*r*)^ and randomly selects 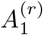 of the pieces from study 1, 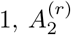 of the pieces from study 2 and so forth. It then combines these selected pieces into a pseudo-study. Since *n*_1_ and *n*_2_ differ, the size of the pieces and thus the number of observations contributed may differ between the studies. The size of a pseudo-study may also differ from the sizes of the observed studies. For example, let *b* = 5, *K* = 3, *n*_1_ = 50, *n*_2_ = 75, *n*_3_ = 150 and the *r*^*th*^ study bag, **A**^(*r*)^ = [1 0 4]^*T*^. We generate the pseudo-study by resampling one fifth of study 1’s 50 observations (i.e., 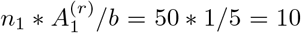 observations) and four fifths of study 3’s 150 observations. Other examples of pseudo-studies generated at different bag sizes are illustrated in Figure 1.

**Fig 1:**
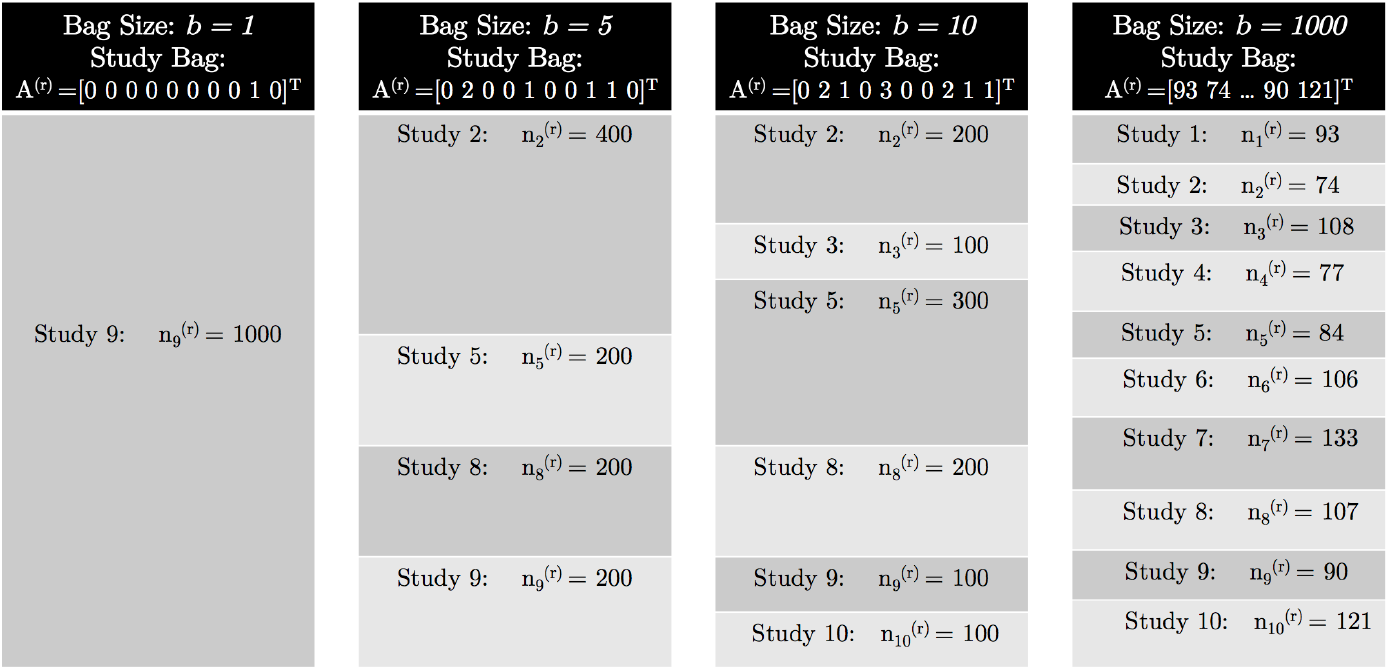
*Illustration of the effect of the bag size tuning parameter b on the resulting pseudo-studies. This example assumes K* = 10, *and observed study sample sizes: n*_1_ = … = *n*_10_ = 1000. *We show four example pseudo-studies. The black rectangles show the bag size, b, and example study bags. The remainder of the column is the study strap replicate, with gray blocks denoting a pseudo-study*. 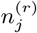 *is the number of observations resampled from the j*^*th*^ *observed study in the r*^*th*^ *pseudo-study*.

We denote the *r*^*th*^ pseudo-study as 𝕊^(*r*)^ and a model trained on the *r*^*th*^ pseudo-study as **Ŷ** ^(*r*)^(·). Similar to how bagging is used in single-study statistical learning (Breiman, 1996a), the study strap can be used to create an ensemble learner, termed *Study Strap Ensemble* (SSE). A full algorithmic description is included in the Supplement (Algorithm 2).

One constraint of the study strap is that observed studies with larger sample sizes will contribute, on average, more observations to pseudo-studies, because observations are resampled in proportion to study sample size. One may instead wish to resample equal numbers of observations from the *k*^*th*^ and *l*^*th*^ observed studies (conditional on 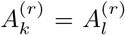) even if the *k*^*th*^ study has a larger sample size. To address this, we introduce the *generalized study strap* resampling scheme, which augments the basic study strap with additional parameters, **N**^*^_*K*×1_. Whereas the standard study strap involves drawing `round` 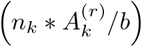 observations from the *k*^*th*^ observed study when constructing a pseudo-study, the generalized study strap resamples `round` 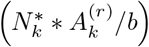. This allows control over both the number of observations contributed by each of the observed studies to a pseudo-study and the induced sample size of a pseudo-study. We provide a detailed algorithmic description in the Supplement (Algorithm 3).

While one could use other resampling schemes such as standard bagging or the randomized cluster bootstrap in multi-study ensemble learning, they do not offer as fine control over the composition of a pseudo-study. Standard bagging resamples in a manner that ignores study membership, thereby providing no way to control how many studies contribute observations to each pseudo-study. Although the randomized cluster bootstrap incorporates the multi-study structure of the problem via its hierarchical procedure, the resampling scheme is restrictive: it resamples with a fixed bag size, *b* = *K*, and draws *A*_*k*_*n*_*k*_ observations from the selected studies. The study strap, on the other hand, affords greater control over the degree of between pseudo-study heterogeneity through the bag size, *b* and the generalized study strap sample size parameters allows one to adjust the size of the pseudo-study. Indeed, the generalized study strap resampling scheme can be viewed as a generalization of the randomized cluster bootstrap method described above (a connection proven below). This distinction is crucial since, as shown later, the performance of an ensemble fit on a study strap replicate can be enhanced substantially by tuning the bag size.

### 2.4. Analytical Results

We present analytical results to demonstrate that the study strap resampling scheme provides a flexible and useful multi-study prediction framework that encompasses earlier methods. We specifically show that the merged dataset (used in the TOM approach), standard (non-hierarchical) bagging, the set of observed studies (used in OSE) and the randomized cluster bootstrap are special cases of the study strap, that arise from specific values of *b* and **N**^*^. The proofs are in Supplementary Section A.2.

#### Proposition 1.

*If the bag size b* = 1 *and sampling is done without replacement, then the set of observed studies (used in OSE) is drawn as the study strap replicate with probability 1*.

#### Proposition 2.

*In the generalized study strap, let the bag size b* = *K, sample size parameters* 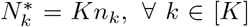, *and assume sampling without replacement. Then the merged dataset (used in TOM) is drawn as the study strap replicate with probability 1*.

Proposition 2 demonstrates how to generate the merged dataset within the generalized study strap resampling procedure. A further and more general connection between the merged dataset and the standard study strap arises when 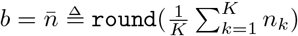. This is discussed next.

#### Proposition 3.

*The standard study strap with* 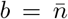 *and sampling without replacement is approximately a delete* 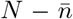 *jackknife bagging of the merged dataset*.

#### Proposition 4.

*Let n* = *n*_1_ = … = *n*_*K*_. *The generalized study strap with b* = *N*, 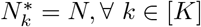 *and sampling with replacement is a non-hierarchical bagging of the merged dataset*.

#### Proposition 5.

*The generalized study strap with* 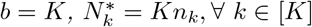 *and sampling with replacement is the randomized cluster bootstrap*.

Taken together, these results show how the parameter *b*, varying between 1 and *N*, provides a tuning mechanism that generates a spectrum of resampling schemes. When *b* = 1 the study strap generates each pseudo-study by resampling from a single study. As *b* grows large (e.g., when 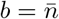), the study strap increasingly ignores study membership and resamples like a standard (non-hierarchical) bagging of the the merged dataset.

### 2.5. Covariate Profile Similarity Weighting

We now focus on “customized training,” when the covariates of the test set are available at the time of model building. “Customized training” has been proposed in both single-study (Kouw and Loog, 2019; Powers, Hastie and Tibshirani, 2015) and multi-study settings (Sun, Shi and Wu, 2015). For example, “multi-source domain adaptation” contains a rich literature on combining datasets in a manner aimed at tailoring prediction to a target dataset. We highlight similarities and differences after describing our method in more detail.

We implemented an approach we refer to as “Covariate Profile Similarity” (CPS) weighting, a weighting scheme that upweights models trained on datasets with feature profiles that are similar to that of the target study, 𝕏_*K*+1_. This approach is motivated by the observation, in the FSCV literature, that combining multiple studies exhibiting covariate distributions similar to that of the test set substantially improves predictive performance and model generalizability. Past work (Kishida et al., 2016) has reported using a K-means clustering approach to select a subset of observed studies (based upon covariate similarity) to merge into one dataset and fit a single model. Here we propose a weighting scheme that can be used in an ensembling setting to leverage information from all available studies.

For a given study or pseudo-study replicate *k*, we define a similarity metric, *s*_*k*_ = 𝒮 (𝕏_*K*+1_, 𝕏_*k*_) ∈ ℝ^+^ such that larger values correspond to greater similarity. This concept is inspired by Minkowski distances and stress functions utilized in the multidimensional scaling literature (Kruskal, 1964a,b; Shepard, 1962). An example metric may simply be the inverse of the *ℓ*_2_ distance between the sample means of the covariates in the two studies considered: 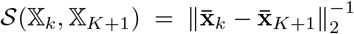. The similarity metric can then be used to adjust the weights for the predictions from the *k*^*th*^ study to obtain final weights 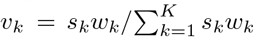. We provide a detailed algorithmic description in the Supplement (Algorithm 1).

### 2.6. Ensembling with the Covariate-Matched Study Strap

While generating pseudo-study replicates has the potential to create powerful training sets, it may also require a large number of pseudo-studies before observed studies with high similarity to the target are well-represented. To address this issue, we developed an alternative adaptive approach to embedding covariate similarity into the study strap framework. This algorithm generates pseudo-study replicates via a study strap, but only trains models on pseudo-studies that have a covariate profile similar to that of the target study. This is accomplished through an accept/reject step where the threshold for covariate similarity is updated to be increasingly selective: each accepted pseudo-study updates the threshold to be the similarity metric of the current pseudo-study. The algorithm runs until a certain number of successive pseudo-studies are generated without acceptance for model training. Since the method is stochastic, we propose iterating through multiple accept/reject paths and ensembling all accepted models. This reduces between-seed variability in performance. We abbreviate this method as “AR” for “Accept/Reject.” See the supplement for a full description (Algorithm 4). As in the SSE, we can weight predictions based upon, for example, stacking or CPS.

Leveraging the marginal distribution of covariates in training and test data is a crucial part of transfer learning, dataset shift and domain adaptation (Kouw and Loog, 2019; Sun, Shi and Wu, 2015). A number of methods propose to align the marginal distribution of the covariates between training and test studies by reweighting *samples* before model training (Kouw and Loog, 2019; Sun et al., 2011). Reweighting is often conducted with techniques such as importance sampling (Shimodaira, 2000) or kernel-based methods (e.g., Kernel Density Estimation (Bickel, Brückner and Scheffer, 2009), Kernel Mean Matching (Huang et al., 2007)). These methods seek to estimate weights for each data point in the training set (or even a weight for each element in the design matrix) and then fit models on the reweighted data. Mansour, Mohri and Rostamizadeh (2009) proposed an ensembling scheme that weights the *predictions* from models that have already been trained. These weights were calculated based on the distribution of the covariates of each training study. Their work was motivated by large datasets where reweighting individual samples could prove computationally intensive. In our application also, reweighting samples would incur prohibitive computational expense as the sample size is about 300,000 and dimension *p* = 1000. Aligning the distribution of covariates in the study strap with many previous methods would require an optimization step to be solved for each pseudo-study. Thus we too propose reweighting predictions based on a measure of covariate similarity between the training and test set, that could be calculated quickly and could also be integrated into our resampling scheme via an accept/reject step. However, our approach differs in that the weights associated with each model are not a function of the covariates of a test data point (i.e., we kept the weights fixed for all observations in a test study) as it is in Mansour, Mohri and Rostamizadeh (2009). This was motivated by the observation that the magnitude of heterogeneity of the covariates within a study pales in comparison to the heterogeneity across studies. As a result we expect that the profile of weights would be very similar across observations within the test study.

## 3. Simulations

We sought to design our simulation experiments to emulate characteristics of the data from our motivating application. For example, the datasets exhibited heterogeneity in the conditional distribution of the outcome given the covariates (Supplemental Figure S.7) and heterogeneity in the distribution of the covariates. Importantly, we observed a number of clusters of studies that exhibited similar covariate profiles (Figure 4) and conditional distributions (Supplemental Figure S.7). For example, we noticed multiple groups (clusters) of studies within which the means of the covariates were very similar. The studies that exhibited similar covariate distributions also tended to have similar profiles of marginal covariate correlations (i.e., univariate correlations between each covariate and the outcome). We therefore simulated datasets to test the impact of: 1) between-study heterogeneity in *f* (𝕏_*k*_), 2) between-study heterogeneity in *f* (**y**_*k*_ | 𝕏_*k*_), and 3) study clusters across which (1) and (2) vary. We illustrate the simulation framework in Supplemental Figure S.1a.

### 3.1. Design of Simulation Experiments

We conducted simulations with 100 iterations, each with *K* = 16 training studies (observed studies) and one test study, under various conditions. For each iteration, we fit different methods and estimated out-of-study prediction performance using the root mean squared error (RMSE) on the test study. We define 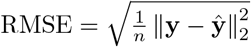, where **ŷ** ∈ ℝ^*n*^ are predicted values. Below we represent each iteration as a single data point in the box plots.

The sample size of each observed study is *n*_*k*_ = 400 for all studies. We included 20 covariates (*p* = 20). These choices mimic the ratio of *p/n*_*k*_ in the neuroscience data where *p/n*_*k*_ ≈ 1000*/*20000. We used smaller *p* and *n*_*k*_ in the simulations because running simulation experiments with the dataset size of the neuroscience data would be prohibitively computationally intensive. Our outcome followed 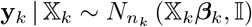. We set 10 of the true model coefficients to be exactly 0, so as to simulate the sparse nature of the neuroscience data. We randomly generated the *p*^*th*^ nonzero model coefficient from the *k*^*th*^ study as 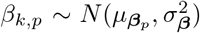 to induce between-study heterogeneity in *f* (**y**_*k*_ | 𝕏_*k*_), where 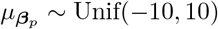. To explore the impact of varying degrees of between-study heterogeneity in the true model, we simulated datasets where the variance 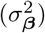 of the distribution from which we drew the true model coefficients ranged across four levels: 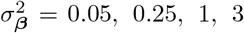. We chose our range to include both favorable and less favorable scenarios for the TOM algorithm, which serves as the reference. The performance of the TOM algorithm ranged from nearly perfect (RMSE is bounded below by 1 since 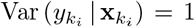, to roughy 150 times that, suggesting that we include a wide range of scenarios. We describe further parameter choices in Supplementary Section B.1.

To induce heterogeneity in *f* (𝕏_*k*_), we randomly drew the overall mean of covariate *p* as 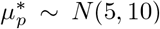, and then drew study specific covariate means 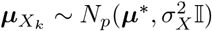 so that 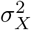 affects the degree of between-study heterogeneity in the means of the covariates. The *i*^*th*^ observation in the *k*^*th*^ study is distributed as, 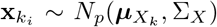 where Σ_*X*_ is a randomly generated covariance matrix that was held constant across studies, but varied between iterations. Levels of 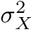 were selected to mimic the ratio of between-study to within-study variability in the covariates that we estimated in the neuroscience dataset. We describe this further in Supplementary Section B.1.

We modeled clusters as groups of studies that shared similar covariate distributions and conditional distributions of *f* (**y**_*k*_ | 𝕏_*k*_). The true model coefficients varied across studies within a cluster by a degree proportional to the between-study variability in model coefficients. If 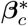 is a baseline vector of true model coefficients for studies in the *c*^*th*^ cluster, then the true model coefficients for the *j*^*th*^ study in the *c*^*th*^ cluster is 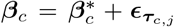 where 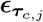 is drawn from 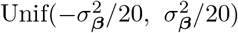. This within-cluster variation was added to ensure that studies within a cluster did not have the same model coefficients or covariate distributions. We simulated study sets with either no clusters or four clusters with four studies per cluster. The number of training and test studies was selected to allow for an even studies/cluster ratios, close to the number of studies (*K* = 15) in our application.

### 3.2. Prediction Approaches Considered in Simulation Experiments

We compared performance of methods for 1) constructing training sets and 2) weighting predictions from different models in the ensemble. For 1) we considered the observed-studies ensemble, study strap ensemble, trained on the merged dataset and Accept/Reject algorithms. We applied average, stacking and CPS weights to each approach. We explore performance in general and customized (CPS and AR) prediction tasks. The study strap used in the SSE and AR algorithms sampled observations without replacement.

We opted to use the LASSO as our single-study learner. The LASSO is commonly used in the human FSCV literature (Moran et al., 2018) because it addresses sparsity, a challenge in our case as well. The LASSO can also be useful in other multi-study settings. Given the sparsity of the simulations, we encountered the challenge that a generic similarity measure comparing covariate profile similarity would equally weight all covariates, including the covariates that did not impact the conditional distribution of *f* (**y**_*k*_ | 𝕏_*k*_). To address this issue, we weight the covariate-wise similarity by a function of the corresponding coefficient estimates. Specifically, we used the similarity measure: 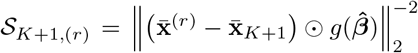, where ⊙ indicates element-wise multiplication (the Hadamard Product) and where the *p*^*th*^ element is

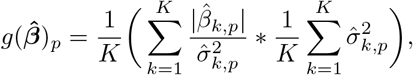

with variance of 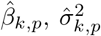, estimated with 500 bootstrap iterations. Our similarity measure was motivated by Sammon mappings and distance measures proposed in multidimensional scaling methods (Sammon, 1969). Our single-study learner sets some coefficients exactly to zero, implying that some of the weights in our similarity measure will also be zero, as desirable.

When tuning the bag size for the SSE and AR algorithms, we used a hold-one-study-out cross validation scheme within the observed studies. We also tuned the LASSO tuning parameter *λ* using a hold-one-study-out cross validation. We kept this tuning parameter fixed across the methods.

### 3.3. Simulation Results

We present simulation results in Figure 2 and Table 1. We provide results in terms of ensembling architecture (OSE, SSE and AR) and weighting schemes: Average (Avg), Covariate Profile Similarity (CPS) and Stacking (STA). We vary three attributes of interest in our simulation: heterogeneity in covariates across studies 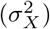, heterogeneity in coefficients across studies 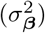, and whether or not there is clustering. Simulation results with a greater range of values for 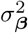 and 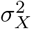 are in Supplementary Section B.2.

**Table 1.** Simulation Results (RMSE / RMSE_TOM_) without clusters (top) and with clusters (bottom). Bold indicates superior performance; no bolded entry in a row indicates that no method was superior to the TOM approach after accounting for Monte Carlo error. Multiple bolded entries indicates (approximate) ties. Monte Carlo error (up to two significant figures) is indicated in parentheses below the corresponding entry. The top row of the methods titles indicates ensembling architecture. The bottom row indicates weighting schemes: Avg (Average), CPS (Covariate Profile Similarity) and STA (Stacking).

| $\sigma_\beta^2 (\sigma_X^2)$ | Observed Studies | | | Study Straps | | | Accept/Reject | | |
| --- | --- | --- | --- | --- | --- | --- | --- | --- | --- |
|  | Avg | CPS | STA | Avg | CPS | STA | Avg | CPS | STA |
| 0.05 (0.0025) | 0.99<br>(0.0034) | 1.01<br>(0.016) | 0.99<br>(0.0026) | 0.99<br>(0.004) | 1.00<br>(0.015) | 1.00<br>(0.0029) | 1.00<br>(0.018) | 1.01<br>(0.03) | 1.00<br>(0.0026) |
| 3.00 | 0.98<br>(0.0057) | 1.01<br>(0.023) | 0.99<br>(0.004) | 0.99<br>(0.0055) | 1.00<br>(0.014) | 1.01<br>(0.009) | 1.01<br>(0.019) | 1.01<br>(0.024) | 1.02<br>(0.0053) |
| 0.05 (400.00) | 3.60<br>(0.81) | 3.01<br>(0.66) | 2.09<br>(0.48) | 6.07<br>(1.1) | 5.78<br>(1) | 0.95<br>(0.14) | 5.01<br>(0.92) | 5.01<br>(0.91) | 4.23<br>(0.9) |
| 3.00 | 1.55<br>(0.25) | 1.42<br>(0.22) | 1.21<br>(0.16) | 2.14<br>(0.27) | 2.05<br>(0.26) | 1.40<br>(0.19) | 1.85<br>(0.32) | 1.83<br>(0.39) | 1.47<br>(0.16) |
| $\sigma_\beta^2 (\sigma_X^2)$ | Observed Studies | | | Study Straps | | | Accept/Reject | | |
|  | Avg | CPS | STA | Avg | CPS | STA | Avg | CPS | STA |
| 0.05 (0.0025) | 1.00<br>(0.0056) | **0.85**<br>(0.017) | 1.00<br>(0.0049) | 1.00<br>(0.0053) | 0.91<br>(0.014) | 1.00<br>(0.004) | 0.89<br>(0.02) | **0.83**<br>(0.025) | 0.99<br>(0.0037) |
| 3.00 | 0.99<br>(0.008) | **0.81**<br>(0.022) | 0.99<br>(0.0063) | 0.99<br>(0.0071) | 0.88<br>(0.015) | 0.99<br>(0.0053) | 0.84<br>(0.022) | **0.77**<br>(0.029) | 1.01<br>(0.0047) |
| 0.05 (400.00) | 5.10<br>(0.58) | **0.28**<br>(0.02) | 1.69<br>(0.19) | 4.65<br>(0.41) | 0.61<br>(0.085) | 0.69<br>(0.14) | 2.48<br>(0.31) | 0.36<br>(0.029) | 1.35<br>(0.19) |
| 3.00 | 8.19<br>(0.7) | **0.47**<br>(0.026) | 2.24<br>(0.26) | 4.71<br>(0.46) | 0.70<br>(0.058) | 0.78<br>(0.056) | 2.47<br>(0.32) | 0.56<br>(0.033) | 1.51<br>(0.17) |

**Table 2.** Relative Mean Squared Error of Prediction Methods in the Voltammetry Data. Entries are RMSE / RMSE_TOM_. Bold indicates best performance among those considered. The top row of the methods titles indicates ensembling architecture. The bottom row indicates weighting schemes: Avg (Average), CPS (Covariate Profile Similarity) and STA (Stacking).

| Covariate Processing | Observed Studies |  |  | Study Straps |  |  | Accept/Reject |  |  |
| --- | --- | --- | --- | --- | --- | --- | --- | --- | --- |
|  | Avg | CPS | Stack | Avg | CPS | Stack | Avg | CPS | Stack |
| Raw | 4.788 | 2.435 | 1.449 | 1.056 | 1.032 | 1.082 | 0.893 | **0.884** | 0.972 |
| Derivative | 2.808 | 1.506 | 1.427 | 1.036 | 1.029 | 1.033 | 0.960 | **0.953** | 0.967 |

**Fig 2:**
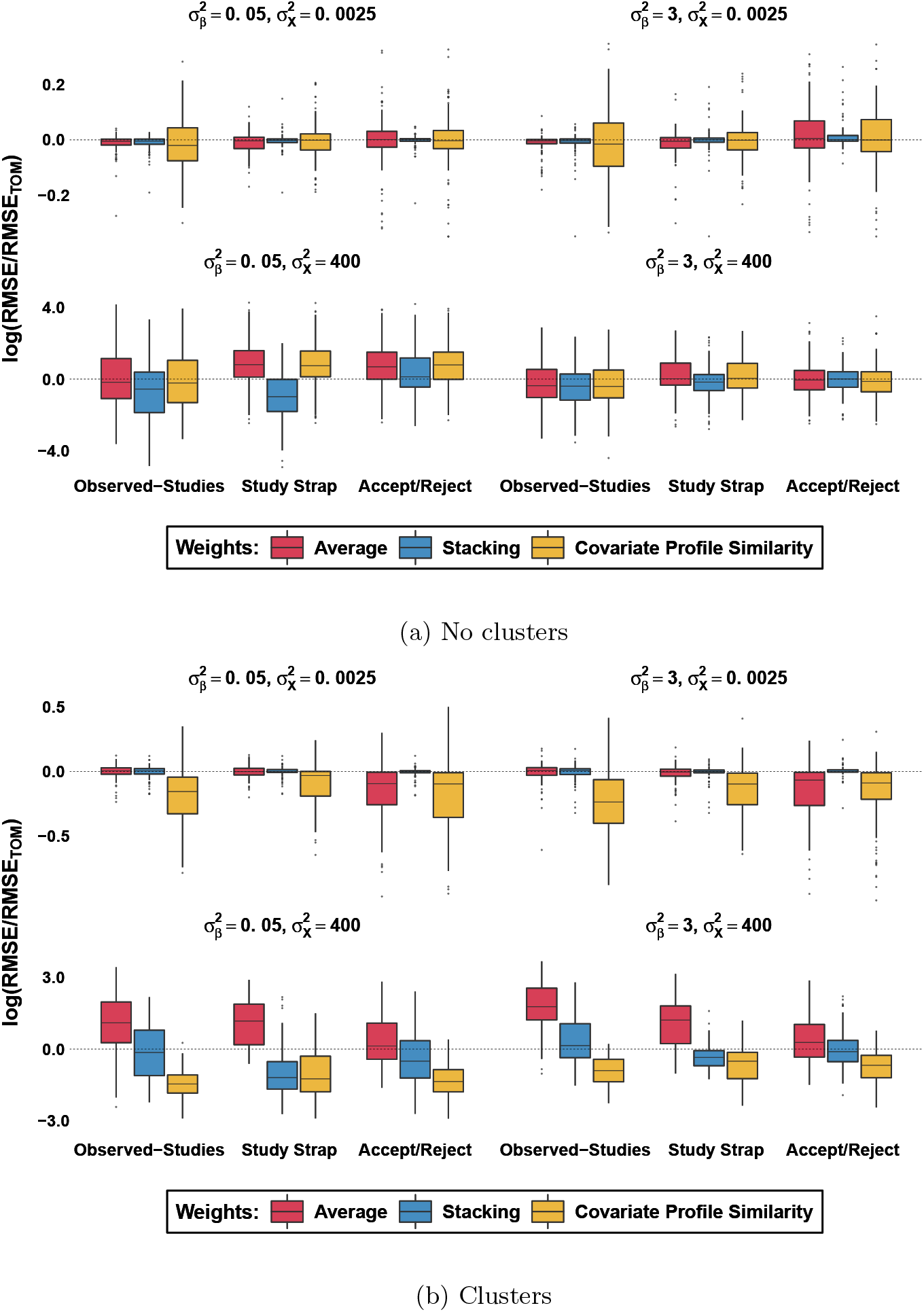
Simulation results for different levels of between-study heterogeneity in true model coefficients, 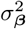 (varies across columns), and distribution of covariates, 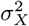 (varies across rows). Each observation in a plot is the log ratio of the out-of-study-RMSE from a single test study (from the corresponding method) to the out-of-study-RMSE of the TOM algorithm (RMSE_TOM_). Each box plot is comprised of 100 iterations.

When heterogeneity in the covariates is low and there is no clustering, ensembling methods tend to perform comparably to merging the data and fitting a single model (i.e., the TOM). As heterogeneity in the covariates rises in the no-clustering case, ensembling methods tend to perform comparatively worse to the TOM (with the exception of SSE with stacking weights).

Our proposed approaches (SSE, AR) yield the largest improvements in performance in settings where there is study clustering. In this setting, CPS weighting with any ensembling scheme confers substantial benefit. Similarly, the AR algorithm improves performance relative to other ensembling methods, but is strongest when paired with CPS weights. The SSE with stacking weights also exhibits considerable gains in performance above non-customized prediction methods. Importantly, the SSE with stacking weights improves performance considerably (even in the no-clustering case) in many of the settings explored and rarely degrades performance. This approach is flexible as it does not require access to the covariates of the test set or specification of a similarity measure.

Importantly, the results emphasize the utility of the hierarchical nature of study strap resampling. We investigated the effect of bag size on performance, graphing the test-set RMSEs against *b* for the AR algorithm (Figure 3a) and for the SSE (Figure 3b; Supplemental Figure S.4). We present results for the AR and SSE algorithms in simulations with clusters to emphasize the utility of bag size tuning. In the AR algorithm, as 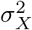 grows, so does the optimal bag size. We emphasize this relationship further in the Supplement (Figure S.5). Correctly tuning the bag size can substantially enhance prediction performance of the algorithm. Similar to the AR, the SSE tended to perform better at lower bag sizes when 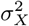 was small. But at these low levels of heterogeneity in the covariates, performance of the SSE varied only moderately as a function of the bag size. Exploring the dependence of the SSE performance on the bag size in these settings required zooming in (i.e., scaling the y-axis), where the impact of mild Monte Carlo error made the bag size curve appear wiggly. The benefit of tuning the bag size for the SSE is, however, very strong at higher levels of 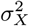. While the optimal bag size with average weights was occasionally in the middle, the benefit of the bag size was most evident when using stacking weights. In the SSE algorithm, the optimal bag size with stacking weights occurred at intermediate values of *b* in simulation settings with larger values of 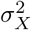. Indeed, the performance at intermediate values of *b* was vastly superior to the extremes. This emphasizes the benefit of the study strap: the study strap ensemble with stacking weights, when using a bag size within a fairly wide neighborhood of the optimal value, substantially outperforms a standard bagging without replacement (equivalent to the SSE with *b* = *n*_*k*_), the OSE (i.e. the SSE with *b* = 1) with stacking weights, or the TOM. The study strap at *b* = *K* = 16 is equivalent to a randomized cluster bootstrap sampled without replacement and with a smaller pseudo-study sample size than a pseudo-study produced by a standard randomized cluster bootstrap. That the performance of the study strap at *b* = *K* is substantially worse than at the optimal bag size suggests that relying on standard hierarchical resampling schemes that use a fixed bag size may be suboptimal for multi-study ensembling. Figure 3b underscores the spectrum between the OSE and the TOM algorithm, generated through varying the bag size *b*. The performance of high bag sizes is nearly identical to that of the TOM algorithm and the low bag sizes approaches that of the OSE (the SSE with *b* = 1) as expected.

**Fig 3:**
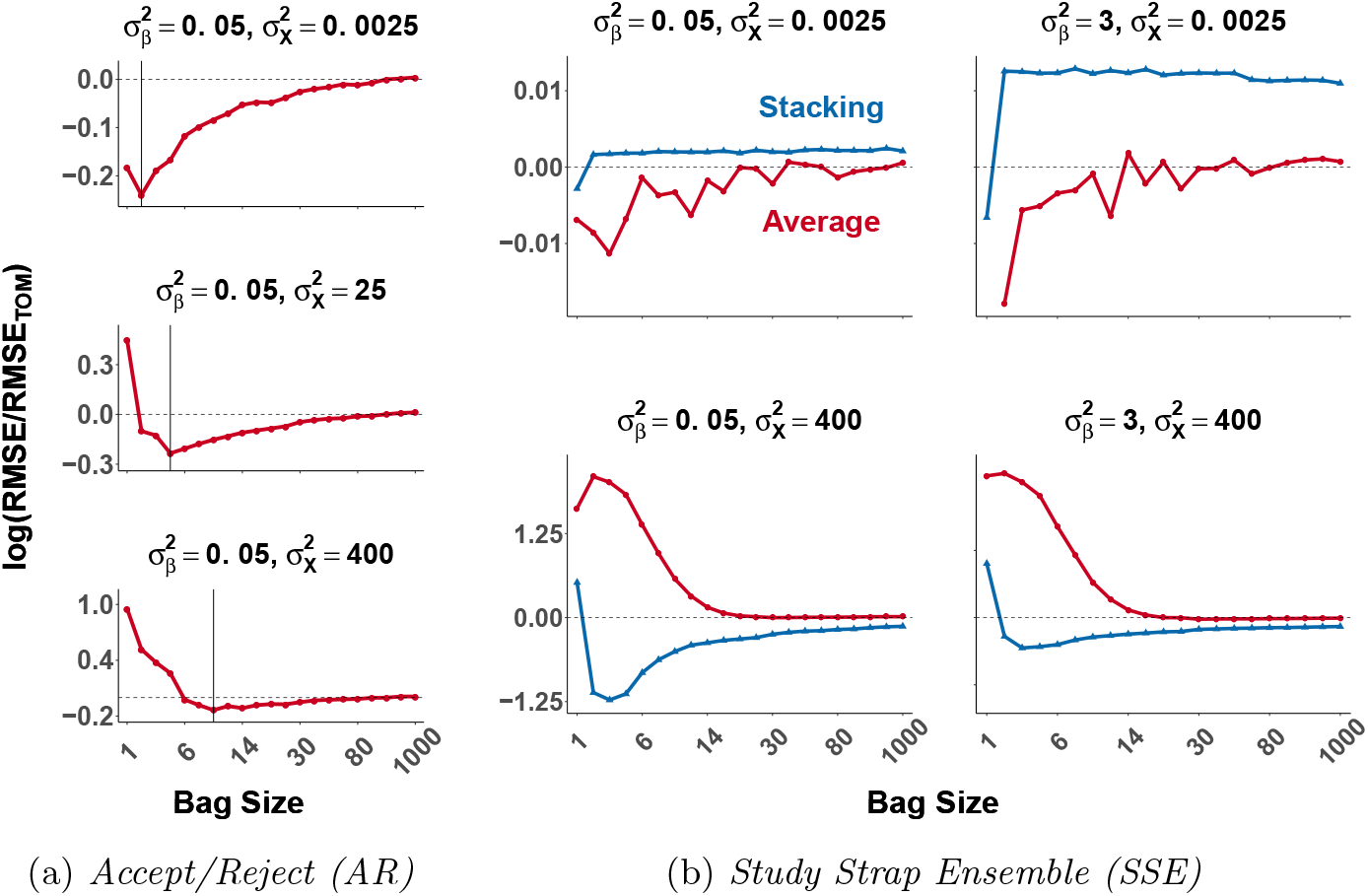
Average test performance as a function of bag size. Points are connected with lines for clarity. The vertical scale is the log of the RMSE in the test study, and divided by the corresponding RMSE of the TOM. Horizontal line indicates performance of the TOM. (a) AR with average weights.Vertical lines indicate the optimal bag size which increases with 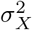. (b) SSE with stacking weights achieves optimal performance at intermediate values of bag size for larger values of 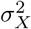.

CPS weights appear to be most effective for lower bag sizes. Indeed, the optimal bag size for the SSE with CPS weights tends to occur at *b* = 1 or close to it (Supplemental Figure S.4). More sophisticated bag size tuning schemes may be an important area of future research: in many cases the tuned bag size for the AR and SSE algorithms were close but not equal to the optimal bag size for out-of-sample testing.

Taken together, these simulations demonstrate that the study strap framework and CPS weights confer substantial benefit particularly when clustering exists in the studies. In cases where our methods do not improve performance, they rarely degrade it compared to standard ensembling methods. Importantly, the simulations broadly display the utility of the bag size.

## 4. Neurochemical Sensing Application

The scientific motivation for our multi-study approach arises from the application of voltammetry to estimate neurotransmitter concentration in awake, behaving human participants (Kishida et al., 2016). We focus on the estimation of dopamine, an important molecule that is thought to underlie learning, reward, and many psychiatric and neurological conditions such as drug addiction and Parkinson’s disease (Volkow, Wise and Baler, 2017). To estimate neurotransmitter concentration, investigators train models on datasets generated in vitro (i.e., where true concentrations are know) and then apply them to measurements made in the brain. Technically, electrodes used to make these measurements (in vitro or in vivo) vary slightly in their construction and slight variations in the experimental setup in which the measurements are made are unavoidable. Thus, each electrode generates a different in vitro dataset, which is representative of a unique observed study in our approach. Our goal was to improve the generalizability of models generated using in vitro data by developing methods that would generate more reliable neurochemical concentration estimates when applied to measurements made in a different context (e.g., the brain). However, no gold standard measure exists to measure neurotransmitters in the brain. Thus to assess model generalizability, we trained models on a subset of studies (i.e., in vitro observed studies, each produced on a different electrode) and evaluated model performance on held out studies (different in vitro observed studies, each produced on a different electrode).

### 4.1. Data Description

The data studied here are from 15 studies (in vitro datasets) each including roughly 20,000 observations. The covariates are electrical measurements (current measured in nanoamps (nA)) collected at 1000 discrete voltage potentials (i.e., the number of covariates, *p* = 1000). The vector of covariates for each observation is called a “Cyclic Voltammogram” (CV) which can also be viewed as a single functional covariate. In each observation, the outcome is a measurement of neurotransmitter concentration in nanomolars (nM). Neuroscientists have analyzed voltammetry data based not only on the raw covariates themselves (Rodeberg et al., 2017) but also on a numerical estimate of the derivative of the raw covariates (the derivative of the current with respect to voltage potential index), in an effort to improve between-study comparability (Bang et al., 2020; Kishida et al., 2016). For this reason, we present results using both. The data are described in greater depth in the Supplement (Section C.1).

### 4.2. Modeling and Methods

We applied our proposed methods to these data using Principal Component Regression (PCR). We found that PCR produced superior cross-study predictive performance compared to regularized regression methods often used in human FSCV (Kishida et al., 2016) (Supplemental Figure S.9). Use of PCR is common in FSCV in rodent experiments (Rodeberg et al., 2017); however, our application of it differs from the standard practice (Keithley and Wightman, 2011; Rodeberg et al., 2017). Kishida et al. (2016) demonstrated that the manner in which PCR is typically applied in rodent FSCV studies (Keithley and Wightman, 2011) results in lower prediction performance than methods currently implemented in human FSCV. Our success with utilizing PCR required larger training datasets and tuning the number of retained principal components using cross-validation, both of which are uncommon practices in the FSCV literature (Keithley and Wightman, 2011; Rodeberg et al., 2017). As a sensitivity analysis, we implemented functional data analytic methods such as functional regression and functional Principal Components Regression (fPCR) using basis splines and found that these exhibited inferior performance in this application. All methods were tested with a hold-one-study-out validation to estimate out-of-study prediction error (RMSE). Since the outcome is chemical concentration, we imposed the non-negativity constraint 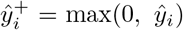 to ensure scientific coherence of the predictions. This only slightly impacted performance.

We selected tuning parameters with a hold-one-study out cross validation scheme. As we also implemented a hold-one-study out procedure to estimate out-of-study prediction performance, we tuned the tuning parameters only within the studies serving as training studies for that fold. To ensure parameter tuning was not responsible for differences in performance between methods, the tuning parameter value was held constant across all of the methods explored (e.g., TOM, AR, OSE).

In designing our similarity measure, we were motivated by the observation that the vector of covariates was high dimensional but the vector of model coefficients appeared sparse (based upon coefficient estimates from regularization methods such as the LASSO). As we describe in greater detail (Supplement Section C.1), our similarity measure was based upon a dimension reduction step where for each study ***ν***_*k*_ = *g* (𝕏_*k*_), ***ν***_*k*_ ∈ ℝ^8^ is a vector summarizing the average covariate profile of the *k*^*th*^ study. We then used as a similarity measure 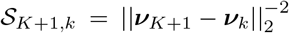 (Supplemental Figure S.8). We used this measure taken on the raw covariates when fitting models on both the original covariates and their numerical derivative.

Given the stochastic nature of our methods, we ran the AR method on 10 separate seeds (each on 10 AR paths) and present the results averaged across seeds. We show the variability in performance between seeds in the Supplemental Figure S.10. Even on seeds associated with the poorest overall performance, our methods still show substantial relative improvement. The study strap used in the SSE and AR algorithms sampled observations without replacement.

As the FSCV data were big, memory considerations even on a computing cluster were a challenge. We thus implemented stacking on a subset of the observations for the SSE and AR algorithms because the design matrix of the stacking regression was large. For the bag size tuning curve, we ran all bag sizes on the AR algorithm on a single path and with a subset of data (*n*_*k*_ = 2500). We selected different bag sizes than in the simulations to accommodate the different sample sizes.

### 4.3. Results

Figure 4 summarizes heterogeneity and clustering in the mean covariate profiles of the studies. Figure 5 summarize the main results using the raw and differentiated covariates. The results demonstrate the strength of the covariate profile similarity weighting scheme. While the observed-studies ensemble, and the weighted versions of it (CPS weights and stacking weights) perform substantially worse than the TOM algorithm, an analysis of the performance of the OSE illustrates the benefits of CPS weights. CPS weighting of the OSE algorithm produced substantial improvements compared to simple average weights. The CPS weights produced a 53.9% and 60.7% reduction in RMSE relative to OSE with simple average weights for the raw and derivative respectively. While weighting the OSE with CPS weights is inferior to using stacking weights, the similarity measure associated with CPS weights allows for the integration of an accept/reject step within the study strap framework. Indeed, the AR algorithm exhibited the strongest performance among all the methods examined. This demonstrates that even when the TOM is superior to existing ensembling approaches used in multi-study learning (i.e., OSE with average or stacking weights), our novel ensembling approaches can substantially out-compete TOM in some scenarios.

**Fig 4:**
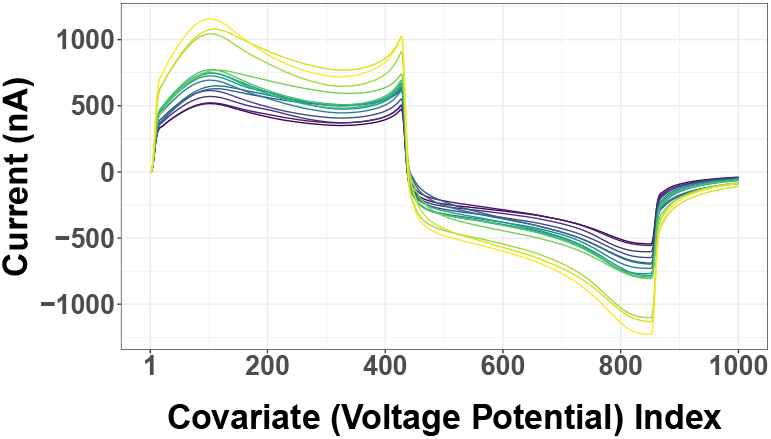
Average covariate profiles. Each curve corresponds to a study (electrode) and is colored by the overall average current. Studies exhibit both variation and clustering in average current.

**Fig 5:**
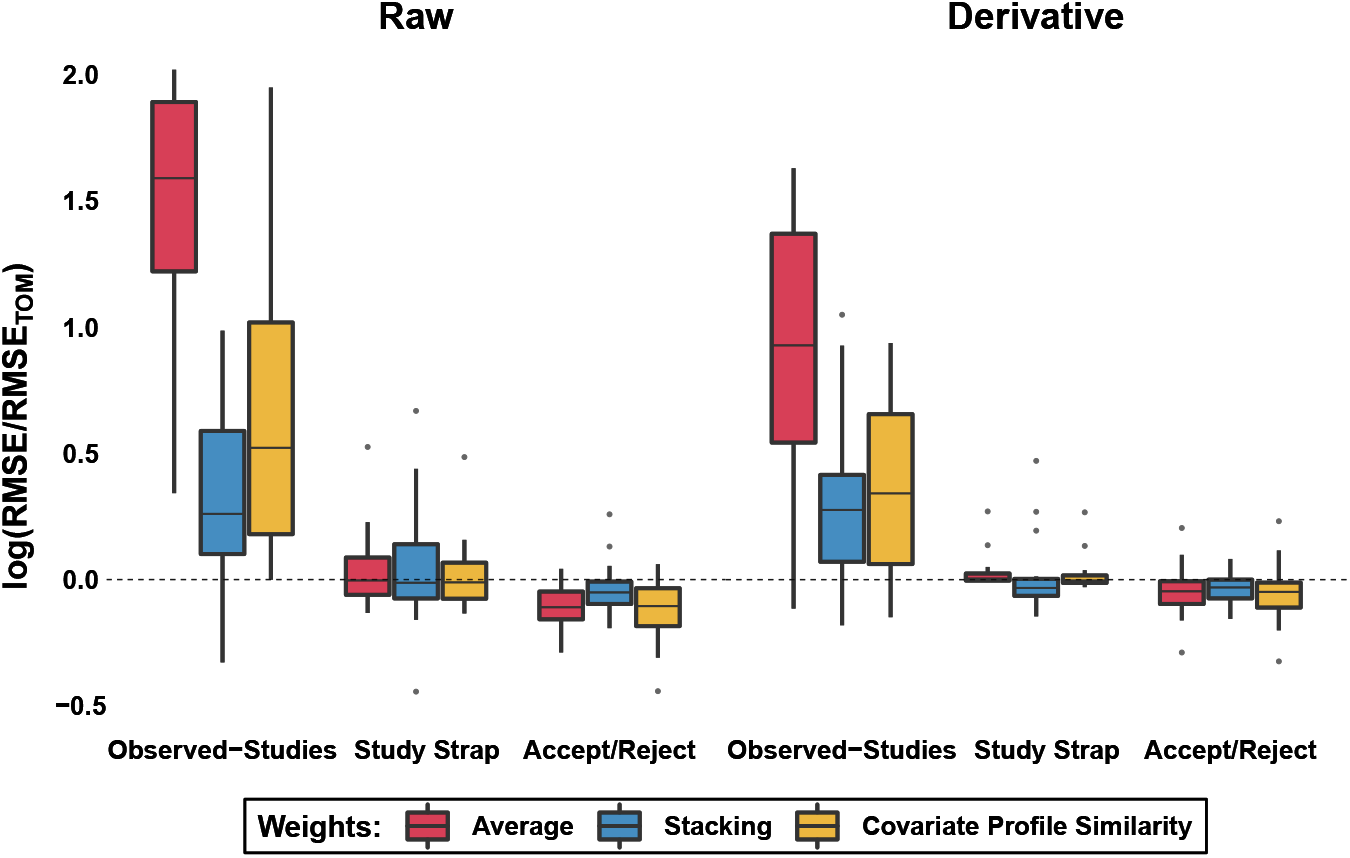
Predictive performance of methods on data using raw covariates (left) and the derivative (right). Dotted line indicates relative performance of the TOM algorithm (Mean raw: RMSE_TOM_ = 416.53; Mean derivative: RMSETOM = 359.64).

Importantly, the use of differentiated covariates shifts the optimal bag size of the AR algorithm with average weights towards higher bag sizes. As apparent in Figure 6, the optimal bag size lies, on average, between roughly 12 and 24 for the raw data, and between 35 and 85 for the derivative. Interestingly, a shift in the optimal bag size for the SSE algorithm with stacking and CPS (but not average) weights also shifts in a similar manner after differentiation (Supplemental Figure S.11). Differentiation modifies the distribution and the between-study heterogeneity of the covariates (Supplemental Figure S.6), likely reducing heterogeneity. The optimal bag size to accommodate the heterogeneity also shifts, demonstrating the utility of the concept.

**Fig 6:**
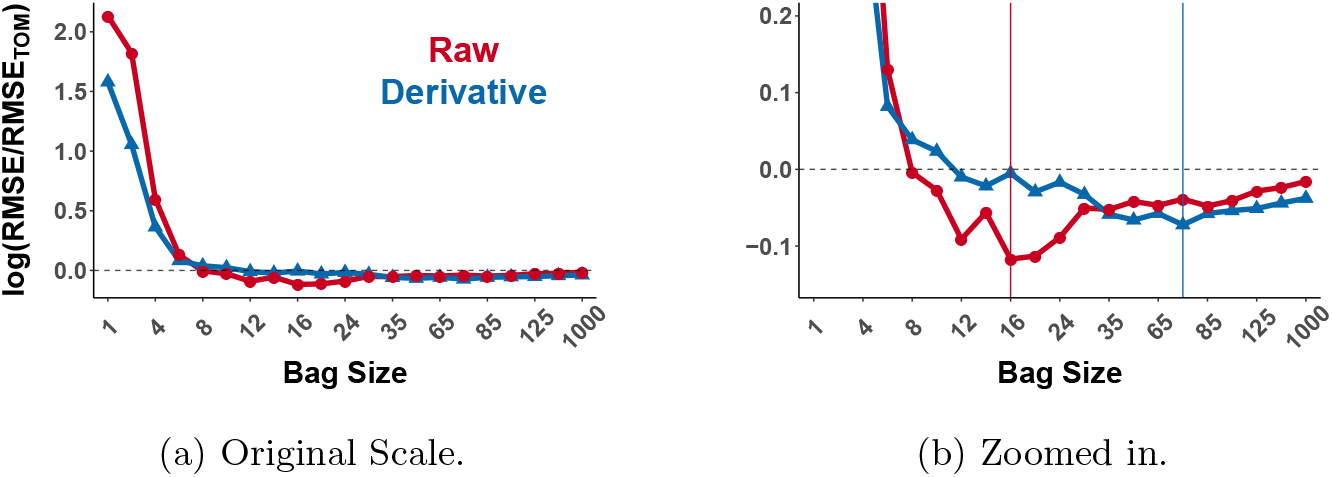
Average Accept/Reject performance on test set as a function of bag size. Derivative shifts the optimal bag size to higher values. Vertical lines indicate optimal bag size. RMSEs are standardized to the RMSE of the T OM algorithm.

While the AR algorithm improved performance by 11.6% relative to the TOM approach when applied to the raw covariates, this number fell to 4.7% when using the derivative. This is consistent with the shift in the optimal bag size associated with using the derivative and echoes a similar pattern in the simulations: as the bag size increases, the benefit of the SSE or AR algorithms, as well as the CPS and stacking weighting schemes, diminishes (Supplemental Figures S.3 and S.4). Relative to the raw covariates, using the derivative improved the performance of all approaches and almost uniformly reduced the difference in performance between any two methods.

## 5. Discussion

We introduce a generalization of multi-study ensemble learning, the study strap, a hierarchical resampling scheme to flexibly generate pseudo-study replicates and to ensemble models trained on these rather than the observed studies. We complement this method with a weighting scheme that incorporates covariate similarity between training data and target populations. Our simulations and data analysis indicate that these methods are robust in that they perform at least as well as our benchmark. The greatest improvements occur when there is clustering of studies. When that is the case, the observed assignment of units to studies is not the most effective way to capture heterogeneity in the distributions of the data, and one might be better off creating clusters in silico instead of relying entirely on the originally observed study labels.

We observe that the bag size tuning parameter *b* of the study strap can help adapt the learning to varying levels of heterogeneity. Our data application provides an example. Using a numerical estimate of the derivative of the covariates as a preprocessing step shifted the optimal bag size to larger values (towards a non-hierarchical bagging of the merged dataset) compared to the optimal bag size of the raw covariates. Similarly, in simulations we saw a consistent shift in the optimal bag size as we varied the degree of between-study heterogeneity in the covariates. Lastly, the optimal bag size is often far from the bag size required to generate pseudo-studies in a manner similar to the standard (non-hierarchical) and randomized cluster bootstrap.

The CPS-based methods implemented here address many of the concerns that chemometric procedures commonly used for FSCV seek to address. Before generating neurotransmitter concentration estimates, standard FSCV statistical methods aim to detect the presence of covariate shift as a proxy for concept shift (Johnson, Rodeberg and Wightman, 2016). Concept shifts commonly arise in FSCV due to between-electrode differences or “drifts” in the electrochemical properties of electrodes upon exposure to, for example, biological tissue (Johnson, Rodeberg and Wightman, 2016). These drifts are accompanied by a shift in the covariate profile. Investigators have historically utilized hypothesis tests to determine whether *f* (𝕏_*train*_) differ significantly from *f* (𝕏_*test*_). CPS weighting and the AR algorithm seek to address this challenge by upweighting studies (or pseudo-studies) that exhibit similar covariate profiles. Our `studyStrap` software package provides over 20 generic similarity measures for the ensembling methods. Context-specific similarity measures (as used in the FSCV data analysis) may be preferable if subject matter knowledge can guide their development. However, even general measures, such as that used in our simulations, may be helpful in a range of scenarios. The similarity measures used here are far from exhaustive. For example, one could extend our approach to nonlinear methods such as kernel-based measures (Gong et al., 2012; Hu et al., 2020) or stochastic neighborhood embeddings (Xu et al., 2019).

Our results demonstrate that despite the statistical challenges present in FSCV research, substantial improvements in cross-electrode generalizability are feasible. Statistical research into this topic is critical as FSCV in humans is new and has, in some domains, unprecedented capacity to provide insight into brain mechanisms underlying human behavior. Moreover, FSCV highlights an important role for statistics in neuroscience: novel applications of statistical learning were instrumental in its successful implementation in humans. These statistical methods have even enabled researchers to use FSCV to measure multiple neurotransmitters simultaneously, a feat thought not to be possible with this technology, even in rodent models (Bang et al., 2020; Montague and Kishida, 2018; Moran et al., 2018).

The study strap and our approach to covariate weighting may be useful in a range of transfer learning settings. Importantly, the study strap achieved considerable gains in performance both with CPS weights and without them(e.g., the study strap with stacking weights). Thus our work is relevant for both domain adaptation (Farahani et al., 2020) and domain generalization (Wang et al., 2021). The methods proposed here may also be useful in multi-task learning, a related transfer learning area in which one seeks to enhance the performance on each of multiple related tasks (i.e., “domains” or studies) by simultaneously training on all tasks (Zhang and Yang, 2021). We hope the methods here are a contribution to the rich literature on transfer learning and multi-study methods, broadly defined.

We conclude by adressing some limitations of our work. An unsolved challenge is to provide an explicit analytical expression for prediction error (RMSE) as a function of the bag size parameter, *b*. While the bag size appears to help account for heterogeneity in the marginal distribution of the covariates in the simulations as well as the neuroscience data, it is clear that the relationship is neither linear nor monotonic and the shape of the relationship appear to vary between settings. Although the AR algorithm was associated with the greatest improvement in performance in the FSCV data, it is the most computationally intensive. Future work may thus seek to replace the accept/reject step with an optimization procedure to increase the speed of the algorithm.

Future work may exploit the functional nature of the FSCV data explicitly, although we found that the functional methods that we implemented (e.g., fPCR) produced inferior performance to standard PCR. Moreover, neither the present work nor any standard methods in the FSCV field, to our knowledge, account for the time-series nature of the data in estimating neurochemical concentration.

Finally, although we have shown progress in enhancing between-electrode generalizability of the models, we cannot verify the generalizability of models trained on in vitro data when applied to data collected in vivo, as this would require an additional gold standard measure of neurochemical concentration in the human brain for which assays do not currently exist. We hope that by enhancing cross-electrode generalizability, generalization of models to data collected in the brain is also improved.

In summary, the study strap and covariate profile similarity weighting are flexible ensembling and weighting schemes that can improve predictive performance in multi-study settings. We hope the present work will contribute to the generalizability of prediction algorithms in neuroscience and beyond.

### 5.1. Reproducibility

We provide the `studyStrap` package on `CRAN` which implements the previously proposed TOM algorithm, OSE and multi-study stacking. It also implements our proposed methods, the study strap ensemble, CPS weighting and the Covariate-Matched Study Strap.

Code and instructions to reproduce analyses are available at: https://github.com/gloewing/studyStrap_Figures

Data from the neuroscience application is available at: https://osf.io/tb8fx/

## 5.2. Acknowledgements

We thank the reviewers for all comments and suggestions. Their feedback helped us to substantially improve the manuscript.

GCL was supported by the NIH, F31DA052153; T32 AI 007358. GP and PP received support from NSF-DMS1810829. KK received support from the NIH, R01 DA048096; R01 MH121099; R01 NS092701; 5KL2TR00142; WFSOM, Phys/Pharm Neurosurgery.

## Appendix A: METHODS AND PROOFS

### A.1. Algorithms

#### Algorithm 1 Ensembling with Covariate Profile Similarity Weighting

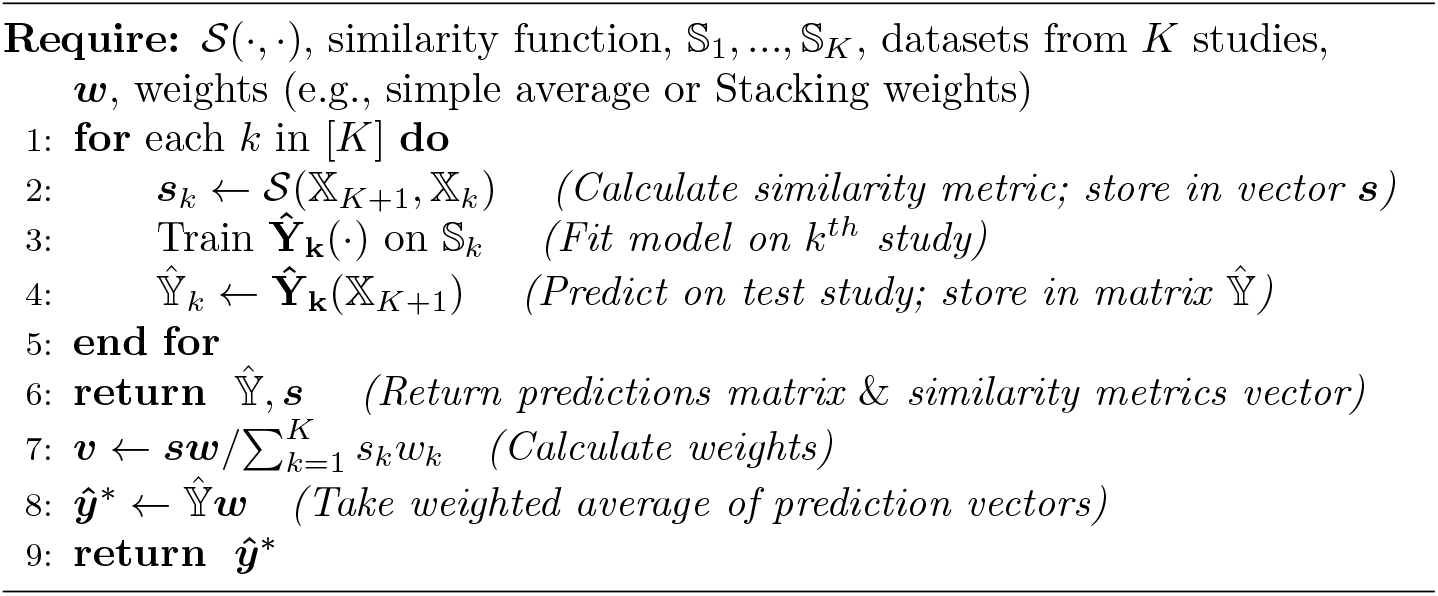

#### Algorithm 2 Ensembling with the Study Strap

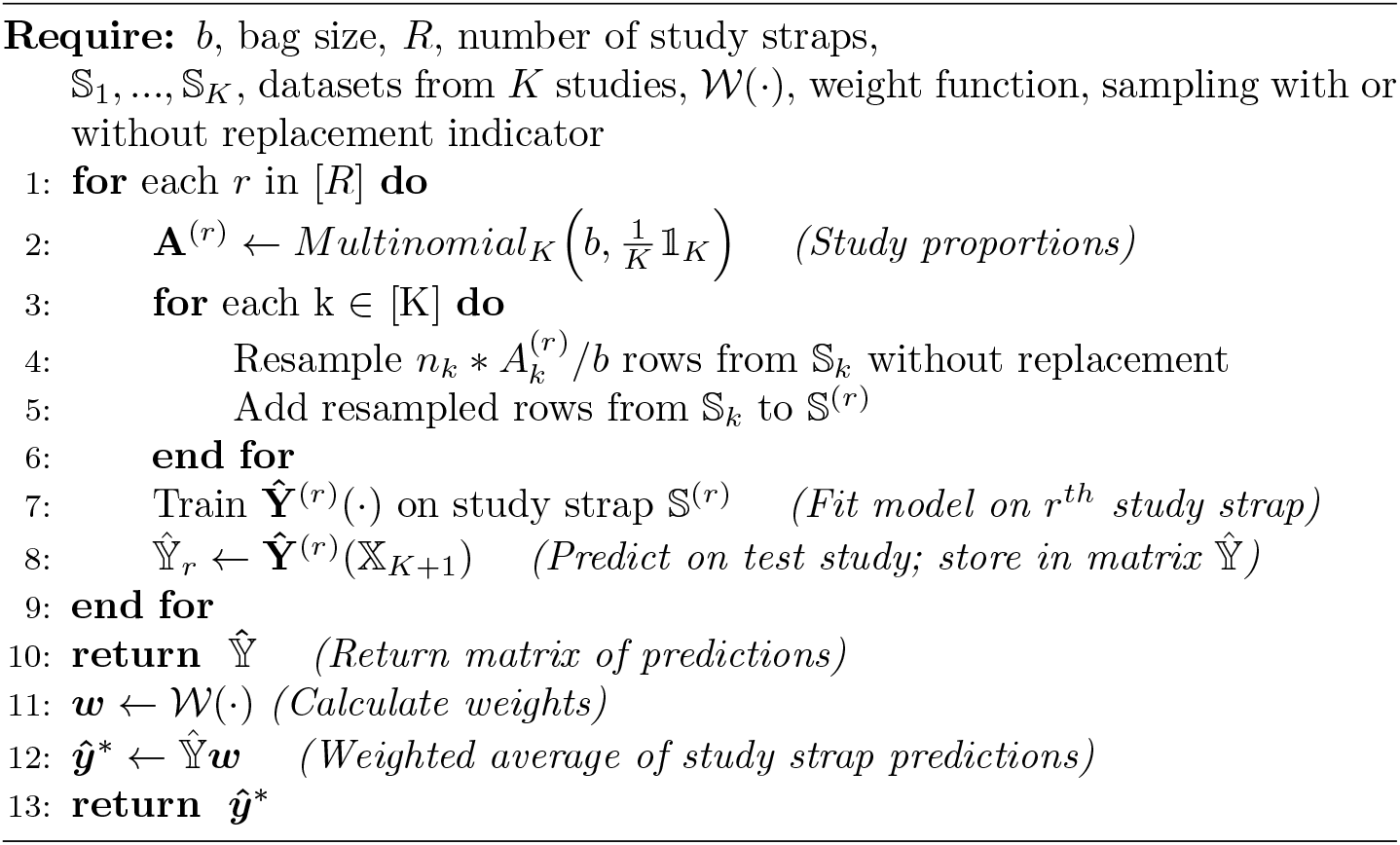

#### Algorithm 3 Ensembling with the Generalized Study Strap

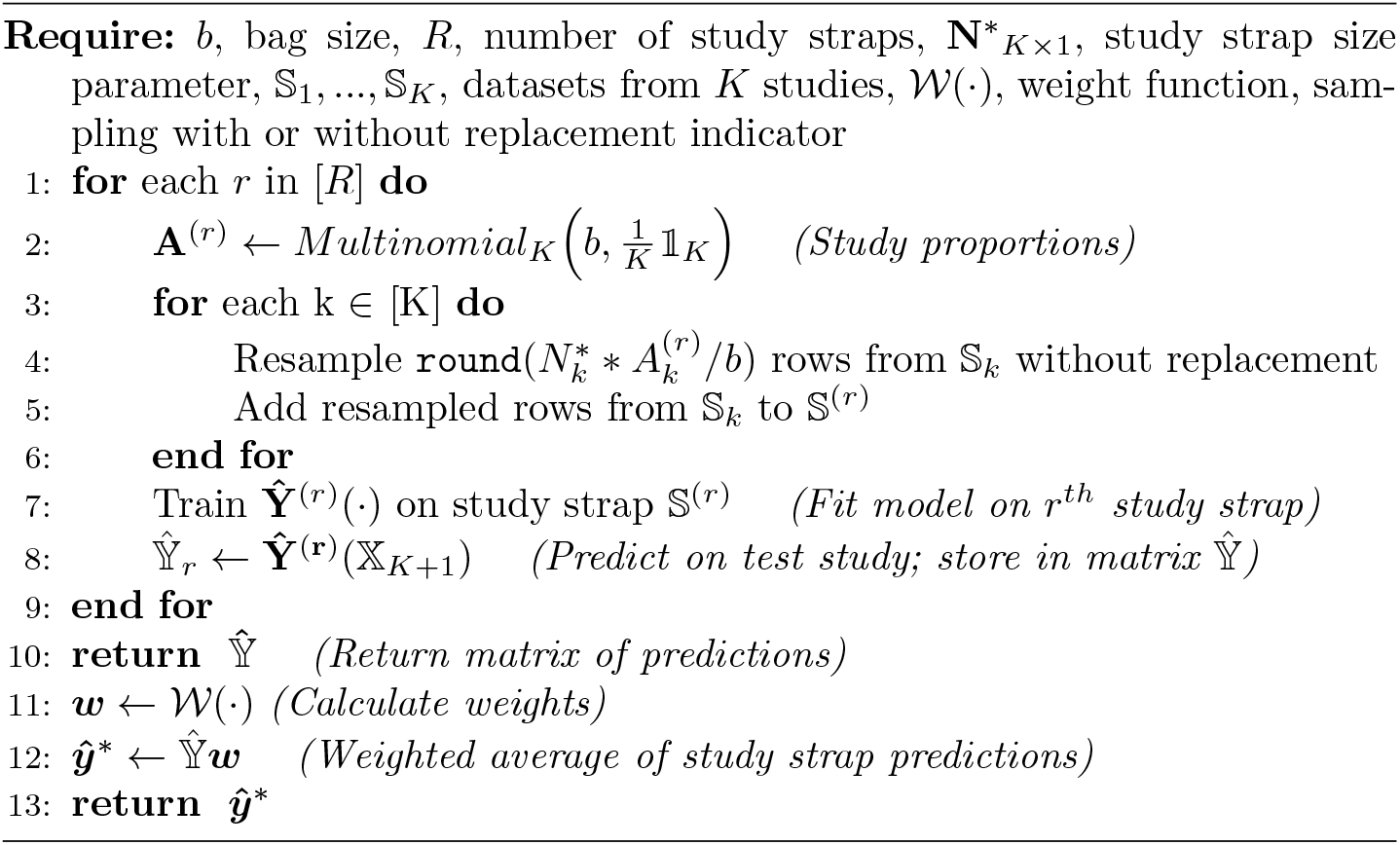

#### Algorithm 4 Ensembling with the Covariate Matched Study Strap (Accept/Reject)

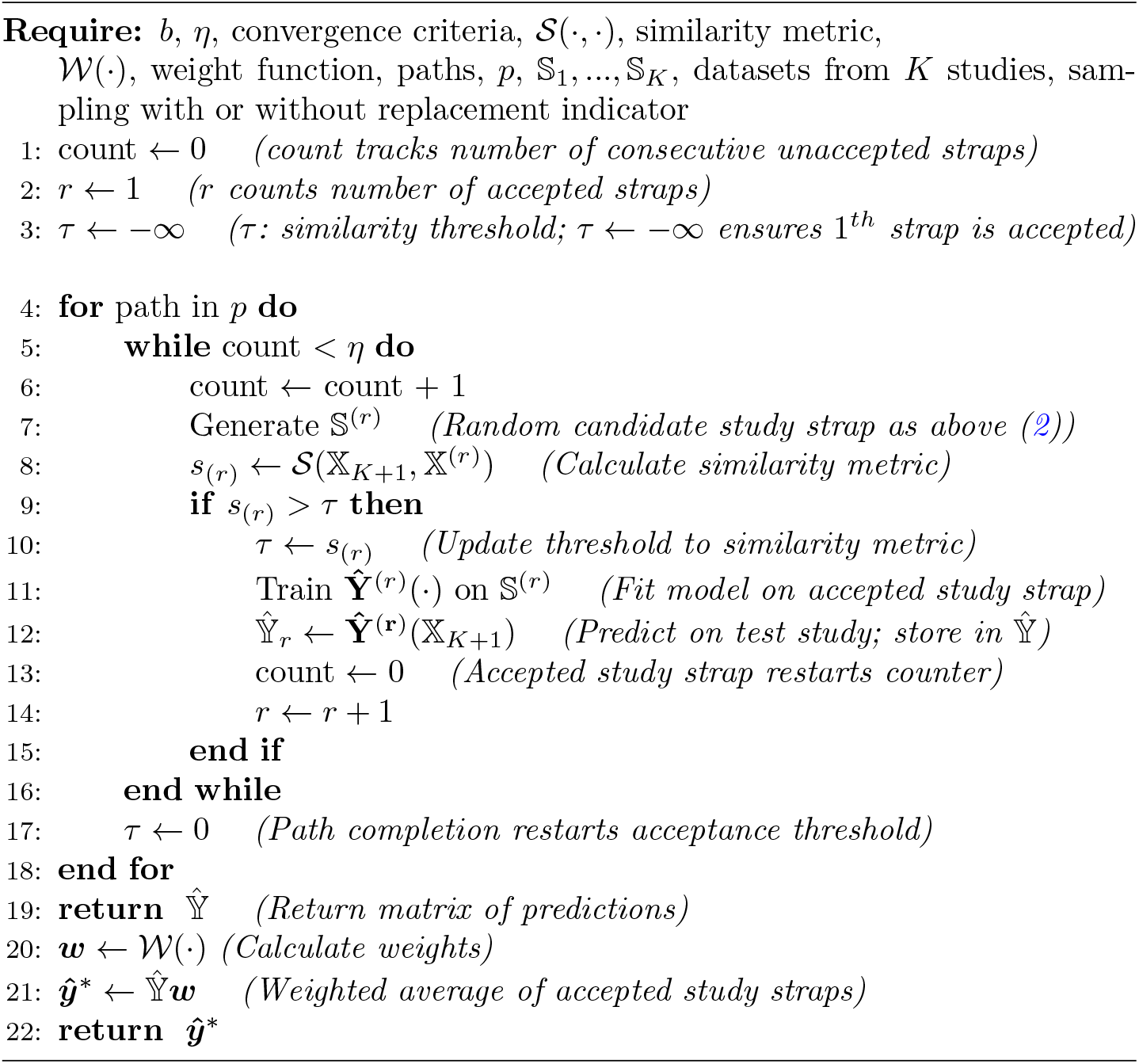

### A.2. Proofs

First, observe that under standard (non-hierarchical) bagging, the number of times an observation is represented in a given bootstrap sample follows the Binomial distribution 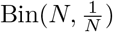 where *N* = Σ_*k*_ *n*_*k*_ is the total sample size. In the study strap, we implement a hierarchical resampling scheme. From the multinomial specification of the vector of counts **A**^(*r*)^, the count 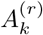 corresponding to the *k*^*th*^ study is *marginally* distributed as a Bin(*b*, 1*/K*).

If one resamples observations with replacement, then the number of times the *i*^*th*^ observation of the *k*^*th*^ study is included in the pseudo-study replicate is distributed as 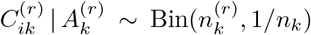 where 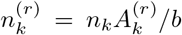. If one resamples without replacement, then 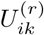 is an indicator that the *i*^*th*^ observation of the *k*^*th*^ study is represented in the *r*^*th*^ pseudo-study replicate, where 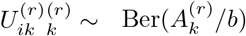.

We point out that under standard (non-hierarchical) delete *N* − *s* jackknife resampling (bagging without replacement) of the merged dataset, 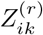, an indicator that the *i*^*th*^ observation of the *k*^*th*^ study is included in the *r*^*th*^ pseudo-study, follows Ber(*s/N*), where *N* = Σ_*k*_ *n*_*k*_ is the total sample size of the merged dataset and *s* is the number of observations resampled from the merged dataset.

#### Proposition 1.

*If the bag size b* = 1 *and sampling is done without replacement, then the set of observed studies (used in the OSE) is drawn as the study strap replicate with probability 1*.

*Proof*. First, observe that for the *r*^*th*^ pseudo-study replicate, all of the observations in 𝕊^(*r*)^ are resampled from a single study, say 𝕊_*k*_, since we draw the *r*^*th*^ study bag from a 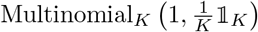. That is, 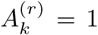 and 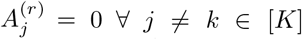. Then 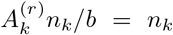 observations are resampled from the *k*^*th*^ study and 0 observations are sampled from all other training studies. When the pseudo-study is resampled without replacement, all observations from the *k*^*th*^ study are resampled so the *r*^*th*^ pseudo-study and the *k*^*th*^ study are identical (i.e., 𝕊^(*r*)^ = 𝕊_*k*_). Since we require each pseudo-study to be unique, (i.e., 𝕊^(*i*)^ ≠ 𝕊^(*j*)^ ∀ *i* ≠ *j* ∈ [*R*]), there are exactly *K* distinct study bags (i.e., *R* = *K*) and corresponding pseudo-studies. Thus, {𝕊_1_, 𝕊_2_, …, 𝕊_*K*_} = {𝕊^(1)^, 𝕊^(2)^, …, 𝕊^(*K*)^}.

It follows directly from the above that that a model fit on a study strap ensemble with *b* = 1 and sampling without replacement produces an ensemble identical to the observed-studies ensemble. That is,

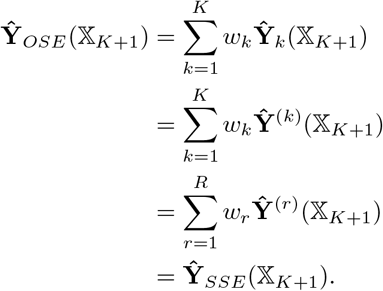

The above proofs stipulate that we cannot have multiple identical pseudo-studies in a study strap replicate in order to achieve an exact equivalence between specific cases of the study strap and the collection of observed studies. Although we feel this stipulation is principled, it is trivial to show that the vector of predictions generated from a study strap ensemble with no limitations on the number of identical models in an ensemble would converge in probability to that produced by the observed-studies ensemble (i.e., as *R* → ∞), through appealing to the Weak Law of Large Numbers:

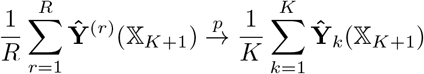

#### Proposition 2.

*In the generalized study strap, let the bag size b* = *K, sample size parameters* 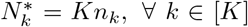, *and assume sampling without replacement. Then the merged dataset (used for TOM) is drawn as the study strap replicate with probability 1*.

*Proof*. Observe that when the pseudo-study is resampled without replacement, this requires that 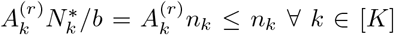 in order to ensure that no observation is resampled more than once. Thus, the only feasible study bag is 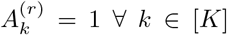 as any other study bag will result in 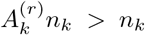 for some *k* ∈ [*K*]. That is, the only possible pseudo-study is constructed by sampling *n*_*k*_ observations without replacement from the *k*^*th*^ study ∀ *k* ∈ [*K*], the merged dataset.

#### Proposition 3.

*The standard study strap with* 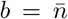 *and sampling without replacement is approximately a delete* 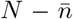 *jackknife bagging of the merged dataset*.

*Proof*. Recall that under standard (non-hierarchical) delete *N* − *s* jacknife resampling of the merged dataset, the indicator 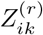 that the *i*^*th*^ observation in the *k*^*th*^ is represented in the *r*^*th*^ pseudo-study is distributed as Ber(*s/N*), where *N* is the total sample size of the merged dataset (i.e., 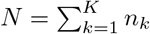), and *s* is the number of observations that we resample (without replacement). Now if we let 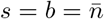, we have 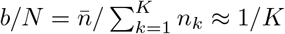, where the approximation is up to the rounding error to ensure an integer bag size (since 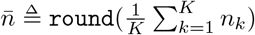). Thus, 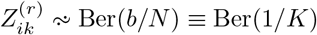.

Now, to demonstrate the stated approximate equivalence, let us show for a study strap with bag size 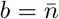 that the indicator variable, 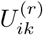, of whether the *i*^*th*^ observation in the *k*^*th*^ study is represented in the *r*^*th*^ pseudo-study is approximately marginally distributed as Ber(1*/K*). First, recall that in generating the *r*^*th*^ pseudo-study, we resample 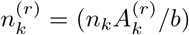 observations from the *k*^*th*^ study, where 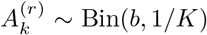. Now the probability that the *i*^*th*^ observation in the *k*^*th*^ study is represented in a given pseudo-study follows approximately the conditional distribution 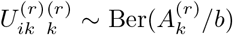, where the approximation is up to the rounding error to ensure an integer sample size for the *r*^*th*^ pseudo-study. Therefore, the marginal probability that the *i*^*th*^ observation in the *k*^*th*^ study is represented in the *r*^*th*^ pseudo-study can be (approximately) expressed as:

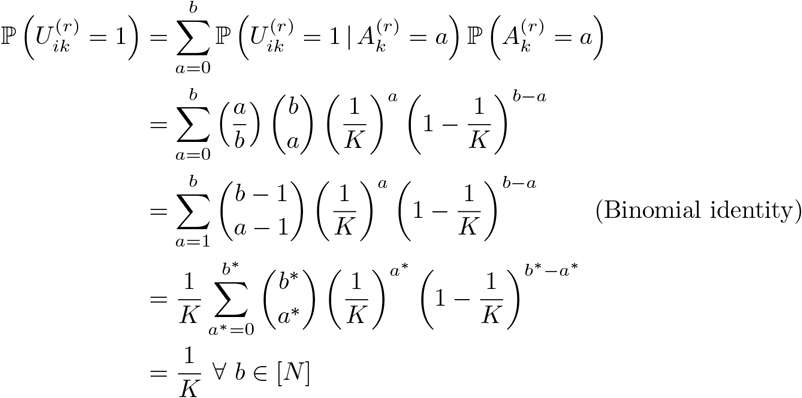

#### Proposition 4.

*Let n* = *n*_1_ = … = *n*_*K*_. *The generalized study strap with b* = *N*, 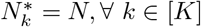 *and sampling with replacement is a non-hierarchical bagging of the merged dataset*.

*Proof*. Recall that under standard (non-hierarchical) bagging of the merged dataset (with replacement), the number of times an observation is represented in a given bootstrap sample follows the distribution 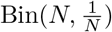 where *N* = Σ_*k*_ *n*_*k*_. In the generalized study strap with the parameters stated above, we implement a hierarchical resampling scheme where the number of times the *i*^*th*^ observation of the *k*^*th*^ study is included in the *r*^*th*^ pseudo-study is 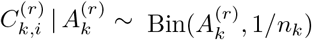 and 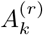 is marginally distributed as Bin(*N*, 1*/K*). Then by the Law of Total Probability, 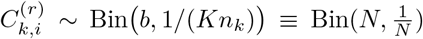, a standard bagging resampling scheme.

#### Proposition 5.

*The generalized study strap with b* = *K*, 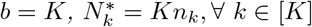 *and sampling with replacement is the randomized cluster bootstrap*.

*Proof*. Recall that in the randomized cluster bootstrap (“Strategy 1”) one samples *K* study labels (with replacement) from which to resample observations. That is, for the primary or study-level sampling step of the *r*^*th*^ pseudo-study, 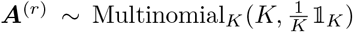. Then, in the secondary or observation-level sampling step, observations are resampled according to the observed study sample sizes selected in the primary step: 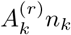 observations are sampled with replacement from the *k*^*th*^ observed study.

Observe that when *b* = *K* and 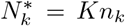, the generalized study strap (re-sampled with replacement) follows the same resampling scheme. Since *b* = *K*, the generalized study strap inherits the primary level resampling step 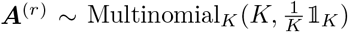. In the observation-level step, *A*_*k*_*n*_*k*_ observations from the *k*^*th*^ observed study are resampled with replacement in the study strap

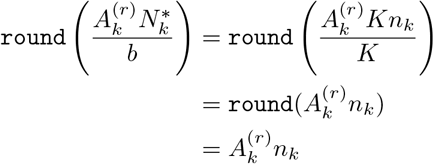

The following distribution can be derived as the *classical occupancy problem* (see Williamson et al. (2009) for a review and proofs), but we include a derivation for completeness.

#### Proposition 6.

*The number of observed studies represented in a given pseudo-study*, 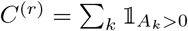, *follows the distribution*, 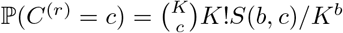

*Proof*. We derive ℙ (*C*^(*r*)^ = *c*). Recall that *C*^(*r*)^ is the number of non-zero counts in the multinomial draw, 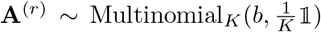 and denote *S*(*b, c*) as Stirling’s number of the second kind (Williamson et al., 2009). The number of non-zero entries of **A**^(*r*)^ is derived by assigning *b* elements into *c* of the total *K* cells since the probabilities are uniform across the cells. There are *K*^*b*^ possible permutations for the multinomial draw and 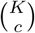 permutations of selecting *c* of the *K* possible cells. Assigning *b* elements into all of the *c* cells (i.e., such that none of the *c* cells are empty) has *K*!*S*(*b, c*) permutations. Thus, the probability that a pseudo-study contains observations resampled from *c* ≤ *K* observed studies is

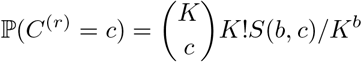

#### Proposition 7.

*The number of observed studies represented in a given pseudo-study*, 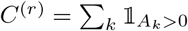, *converges in probability to K as b* → ∞:

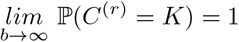

*Proof*.

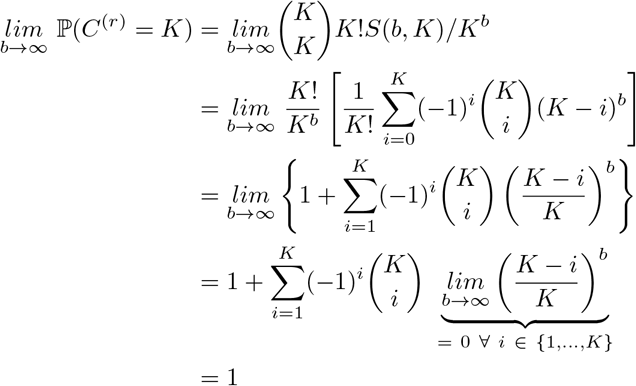

where interchange of the limit and sum follows as *K* is fixed and thus the sum is finite.

### A.3. Stacking Strategy

We used non-negative least squares with an intercept to generate weights. We found that standardizing the weights led to a degradation of performance and so we proceeded without standardizing the coefficient estimates. Thus, the final predictions are:

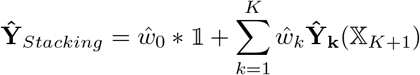

### A.4. Study Strap Stacking Strategy

Below is how we implemented the stacking regression for the study strap ensemble (and AR). This regresses 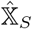 on **y**_*S*_, where

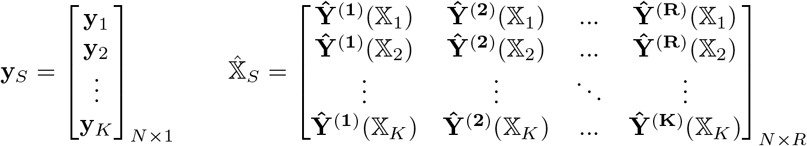

and **Ŷ** ^(**r**)^(𝕏_*k*_) are the predictions on the design matrix of training study *k* using the model trained on the *r*^*th*^ pseudo-study; **y**_*k*_ are the labels from training study *k*. The stacking procedure then proceeds as above.

### A.5. Alternative Study Strap Stacking Strategy

Below is an additional way one could implement the study strap (or AR) analogue of the stacking. This regresses 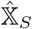 on **y**_*S*_, where

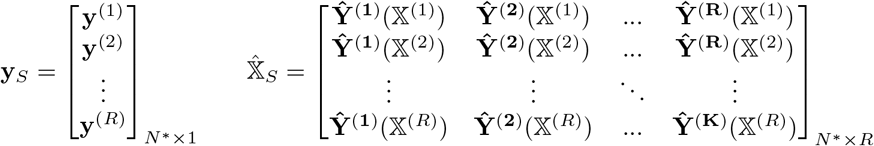

and **Ŷ** ^(**r**)^(𝕏^(*j*)^) are the predictions on the design matrix of the *j*^*th*^ pseudo-study using the model trained on the *r*^*th*^ pseudo-study; **y**^(*j*)^ are the labels from pseudo-study *r*. 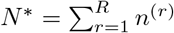 The stacking procedure then proceeds as in standard stacking.

One could similarly construct the stacking regression to weight the OSE or standard Study-Strap Ensemble using the design matrices of the accepted study straps in the AR algorithm (and the models from the OSE or standard OSE respectively). The above variations on standard stacking highlight the flexibility introduced by the study strap framework.

## Appendix B: SIMULATIONS SUPPLEMENTARY MATERIAL

### B.1. Simulation Parameters

Here we describe additional details related to the simulations. When tuning the *λ* tuning parameter for the LASSO, we used a hold-one-study out cross validation scheme and used the same *λ* across studies and methods within a given simulation iteration. In other words, we used the same *λ* for all methods (i.e., TOM, OSE, SSE, AR) to improve comparability across methods. We tested 43 values of *λ* between 0.0001 and 5. When tuning the study strap for the bag size, we fit models to 150 pseudo-studies at each of 21 bag sizes between *b* = 1 and *b* = 1000. We used a hold-one-study-out cross validation scheme to tune the bag size. When generating the study strap bag size performance curve we used 250 pseudo-studies per bag size. When testing the final study strap ensemble with the tuned *b*, we used 500 study straps. We used fewer pseudo-studies during tuning stages to reduce the computational effort (since we were doing a hold-one-study-out cross validation scheme, it was computationally intensive).

When we were tuning the AR bag size, we used a convergence criterion of 10000 consecutive pseudo-studies without an acceptance. We averaged across 3 paths. We used the same parameters for the AR bag size performance curve. For the final AR implementation with the tuned *b*, we used 5 paths and a convergence criteria of 100000 consecutive pseudo-studies without an acceptance. We felt these parameters struck the balance of being computationally feasible and also sufficient for our purposes.

In selecting our degree of between-study heterogeneity in covariates, we tried to mimic the ratio of between-to-within study variance of covariates (averaged across covariates). Specifically, for each covariate, we estimated the variance of covariate means (across studies) divided by the average within-study variance of that covariate. We then averaged this ratio across all 1000 covariates. Formally, we estimated this ratio, denoted as *ϕ*,

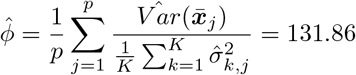

where 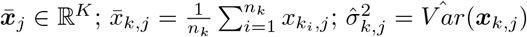 and 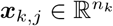.

### B.2. Additional Simulation Figures and Tables

**Table 1.** Simulation Results (RMSE / RMSE_TOM_) without clusters (top) and with clusters (bottom). Bold indicates superior performance; no bolded entry in a row indicates that no method was superior to the TOM approach after accounting for Monte Carlo error. Multiple bolded entries indicates (approximate) ties. Monte Carlo error (up to two significant figures) is indicated in parentheses below the corresponding entry. The top row of the methods titles indicates ensembling architecture. The bottom row indicates weighting schemes: Avg (Average), CPS (Covariate Profile Similarity) and STA (Stacking).

| $\sigma_\beta^2 (\sigma_X^2)$ | Observed Studies | | | Study Straps | | | Accept/Reject | | |
| --- | --- | --- | --- | --- | --- | --- | --- | --- | --- |
|  | Avg | CPS | STA | Avg | CPS | STA | Avg | CPS | STA |
| 0.05 (0.0025) | 0.99<br>(0.0034) | 1.01<br>(0.016) | 0.99<br>(0.0026) | 0.99<br>(0.004) | 1.00<br>(0.015) | 1.00<br>(0.0029) | 1.00<br>(0.018) | 1.01<br>(0.03) | 1.00<br>(0.0026) |
| 0.25 | 0.98<br>(0.0048) | 1.01<br>(0.02) | 0.99<br>(0.0036) | 1.00<br>(0.0046) | 1.02<br>(0.017) | 1.01<br>(0.0086) | 1.01<br>(0.019) | 1.01<br>(0.024) | 1.01<br>(0.0053) |
| 1.00 | 0.98<br>(0.0059) | 1.01<br>(0.022) | 0.99<br>(0.0043) | 0.99<br>(0.0053) | 1.00<br>(0.014) | 1.01<br>(0.0087) | 1.01<br>(0.016) | 1.03<br>(0.023) | 1.01<br>(0.0047) |
| 3.00 | 0.98<br>(0.0057) | 1.01<br>(0.023) | 0.99<br>(0.004) | 0.99<br>(0.0055) | 1.00<br>(0.014) | 1.01<br>(0.009) | 1.01<br>(0.019) | 1.01<br>(0.024) | 1.02<br>(0.0053) |
| 0.05 (25.00) | 1.05<br>(0.092) | 0.99<br>(0.086) | **0.85**<br>(0.073) | 2.20<br>(0.24) | 2.13<br>(0.24) | 0.94<br>(0.073) | 1.99<br>(0.25) | 2.02<br>(0.27) | 1.13<br>(0.09) |
| 0.25 | 1.08<br>(0.11) | 1.04<br>(0.1) | 1.09<br>(0.11) | 1.38<br>(0.13) | 1.37<br>(0.13) | 1.00<br>(0.066) | 1.20<br>(0.13) | 1.22<br>(0.15) | 1.16<br>(0.098) |
| 1.00 | 1.21<br>(0.13) | 1.17<br>(0.12) | 1.13<br>(0.11) | 1.25<br>(0.12) | 1.25<br>(0.12) | 1.06<br>(0.068) | 1.09<br>(0.097) | 1.11<br>(0.099) | 1.11<br>(0.096) |
| 3.00 | 1.36<br>(0.17) | 1.28<br>(0.16) | 1.23<br>(0.14) | 1.25<br>(0.14) | 1.24<br>(0.14) | 1.30<br>(0.14) | 1.16<br>(0.14) | 1.15<br>(0.13) | 1.24<br>(0.12) |
| 0.05 (400.00) | 3.60<br>(0.81) | 3.01<br>(0.66) | 2.09<br>(0.48) | 6.07<br>(1.1) | 5.78<br>(1) | 0.95<br>(0.14) | 5.01<br>(0.92) | 5.01<br>(0.91) | 4.23<br>(0.9) |
| 0.25 | 3.14<br>(0.68) | 2.76<br>(0.62) | 1.66<br>(0.35) | 5.33<br>(1.2) | 5.08<br>(1.2) | 1.63<br>(0.37) | 4.62<br>(0.94) | 4.66<br>(0.98) | 2.27<br>(0.41) |
| 1.00 | 1.84<br>(0.3) | 1.70<br>(0.26) | 1.26<br>(0.19) | 2.97<br>(0.42) | 2.89<br>(0.42) | 1.49<br>(0.23) | 2.82<br>(0.46) | 2.76<br>(0.47) | 2.01<br>(0.39) |
| 3.00 | 1.55<br>(0.25) | 1.42<br>(0.22) | 1.21<br>(0.16) | 2.14<br>(0.27) | 2.05<br>(0.26) | 1.40<br>(0.19) | 1.85<br>(0.32) | 1.83<br>(0.39) | 1.47<br>(0.16) |
| $\sigma_\beta^2 (\sigma_X^2)$ | Observed Studies | | | Study Straps | | | Accept/Reject | | |
|  | Avg | CPS | STA | Avg | CPS | STA | Avg | CPS | STA |
| 0.05 (0.0025) | 1.00<br>(0.0056) | **0.85**<br>(0.017) | 1.00<br>(0.0049) | 1.00<br>(0.0053) | 0.91<br>(0.014) | 1.00<br>(0.004) | 0.89<br>(0.02) | **0.83**<br>(0.025) | 0.99<br>(0.0037) |
| 0.25 | 0.99<br>(0.0074) | **0.82**<br>(0.02) | 0.99<br>(0.0064) | 0.99<br>(0.0069) | 0.90<br>(0.015) | 0.99<br>(0.0058) | 0.88<br>(0.019) | **0.82**<br>(0.028) | 1.00<br>(0.0041) |
| 1.00 | 0.99<br>(0.008) | **0.81**<br>(0.021) | 0.99<br>(0.0066) | 0.99<br>(0.0072) | 0.89<br>(0.016) | 0.99<br>(0.0059) | 0.89<br>(0.015) | **0.83**<br>(0.024) | 1.01<br>(0.0046) |
| 3.00 | 0.99<br>(0.008) | **0.81**<br>(0.022) | 0.99<br>(0.0063) | 0.99<br>(0.0071) | 0.88<br>(0.015) | 0.99<br>(0.0053) | 0.84<br>(0.022) | **0.77**<br>(0.029) | 1.01<br>(0.0047) |
| 0.05 (25.00) | 2.49<br>(0.19) | **0.55**<br>(0.016) | 1.26<br>(0.087) | 2.53<br>(0.23) | 0.66<br>(0.024) | 0.80<br>(0.05) | 1.29<br>(0.091) | 0.64<br>(0.023) | 1.15<br>(0.082) |
| 0.25 | 4.30<br>(0.29) | **0.51**<br>(0.012) | 2.10<br>(0.15) | 2.78<br>(0.23) | 0.67<br>(0.026) | 1.13<br>(0.083) | 1.71<br>(0.15) | 0.61<br>(0.023) | 1.59<br>(0.14) |
| 1.00 (25.00) | 5.82<br>(0.4) | **0.42**<br>(0.012) | 2.84<br>(0.21) | 2.87<br>(0.25) | 0.72<br>(0.049) | 1.34<br>(0.12) | 1.83<br>(0.17) | 0.54<br>(0.022) | 2.00<br>(0.18) |
| 3.00 (25.00) | 5.93<br>(0.42) | **0.43**<br>(0.019) | 2.92<br>(0.23) | 2.60<br>(0.21) | 0.69<br>(0.044) | 1.49<br>(0.13) | 2.01<br>(0.18) | 0.53<br>(0.025) | 2.09<br>(0.18) |
| 0.05 (400.00) | 5.10<br>(0.58) | **0.28**<br>(0.02) | 1.69<br>(0.19) | 4.65<br>(0.41) | 0.61<br>(0.085) | 0.69<br>(0.14) | 2.48<br>(0.31) | 0.36<br>(0.029) | 1.35<br>(0.19) |
| 0.25 | 6.01<br>(0.61) | **0.28**<br>(0.019) | 1.74<br>(0.19) | 4.97<br>(0.44) | 0.61<br>(0.08) | 0.68<br>(0.13) | 2.09<br>(0.23) | 0.42<br>(0.044) | 1.30<br>(0.17) |
| 1.00 | 7.49<br>(0.62) | **0.35**<br>(0.021) | 2.08<br>(0.21) | 5.00<br>(0.44) | 0.62<br>(0.067) | 0.94<br>(0.15) | 2.31<br>(0.26) | 0.45<br>(0.029) | 1.59<br>(0.21) |
| 3.00 | 8.19<br>(0.7) | **0.47**<br>(0.026) | 2.24<br>(0.26) | 4.71<br>(0.46) | 0.70<br>(0.058) | 0.78<br>(0.056) | 2.47<br>(0.32) | 0.56<br>(0.033) | 1.51<br>(0.17) |

**Fig S.1:**
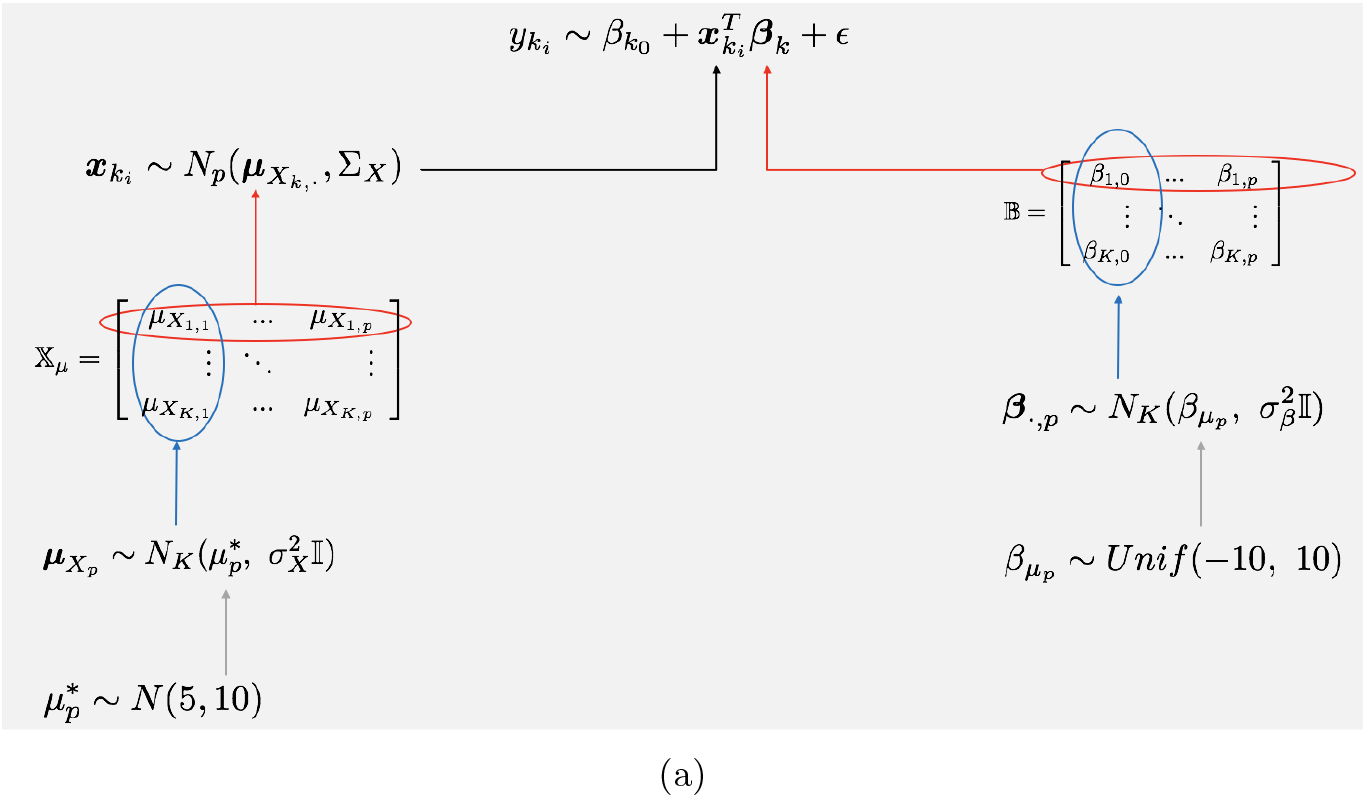
(A) General simulation framework.

**Fig S.2:**
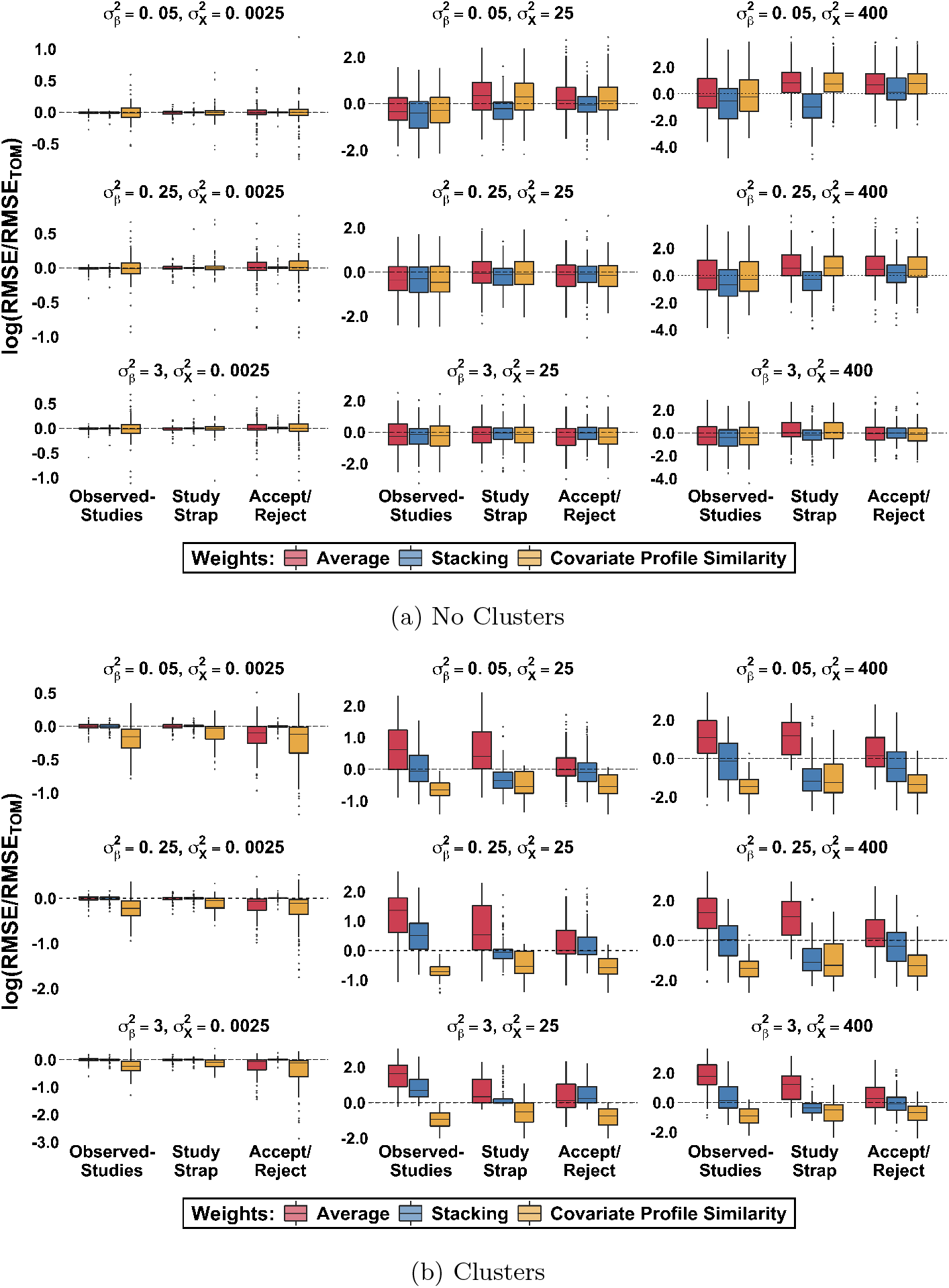
Simulation results on log scale across different cluster sizes and between-study variability (*σ*_***β***_) in ***β***. Each observation in a plot is the out-of-study-RMSE from a single test study (i.e., each whisker plot is comprised of 100 points). *b*^*^ indicates the selected bag size (from tuning) for the study strap ensemble and AR, respectively. Dotted line indicates the relative performance of the TOM algorithm.

**Fig S.3:**
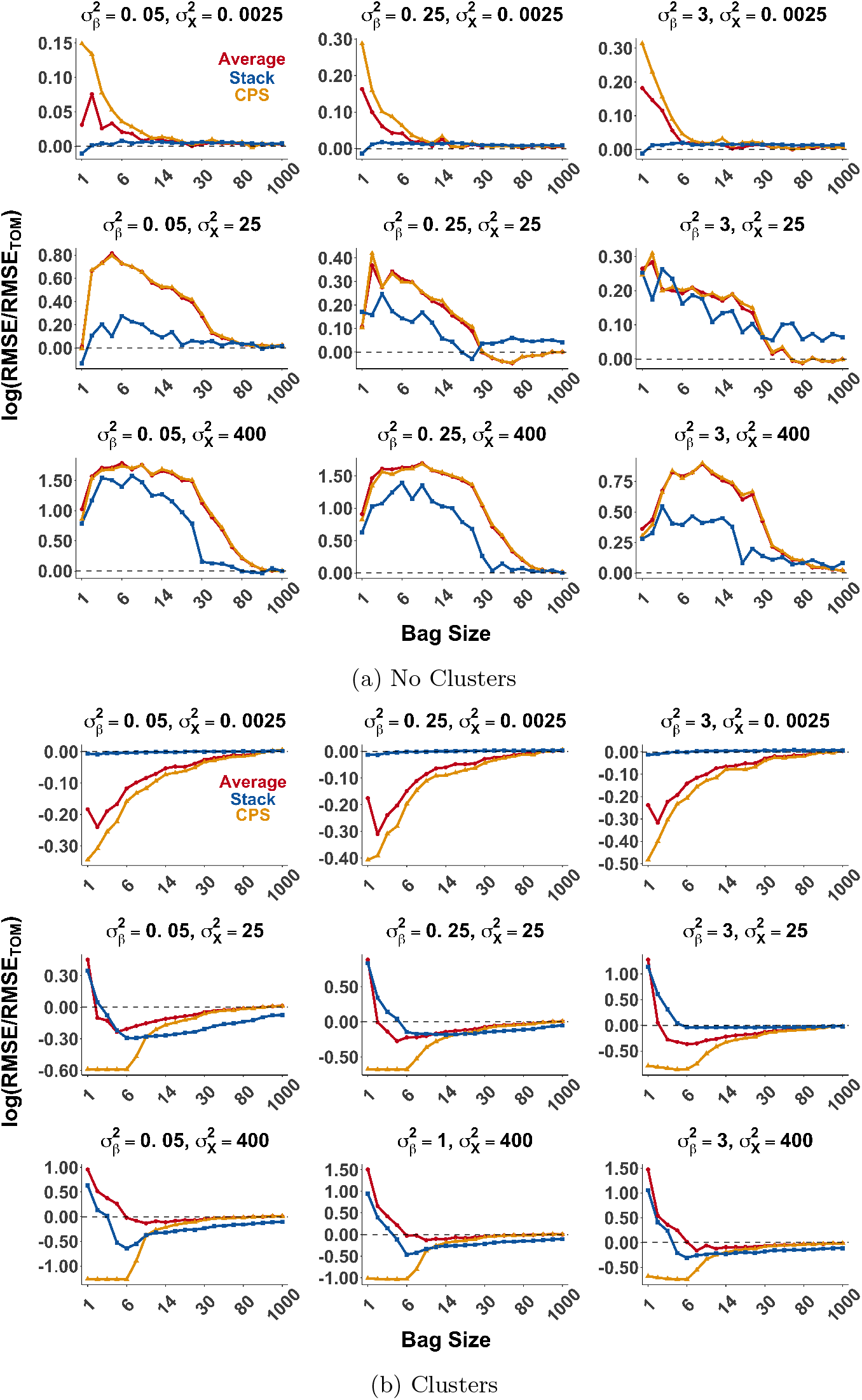
Average performance of AR algorithm on test set as a function of bag size. Relative performance of TOM indicated with black line.

**Fig S.4:**
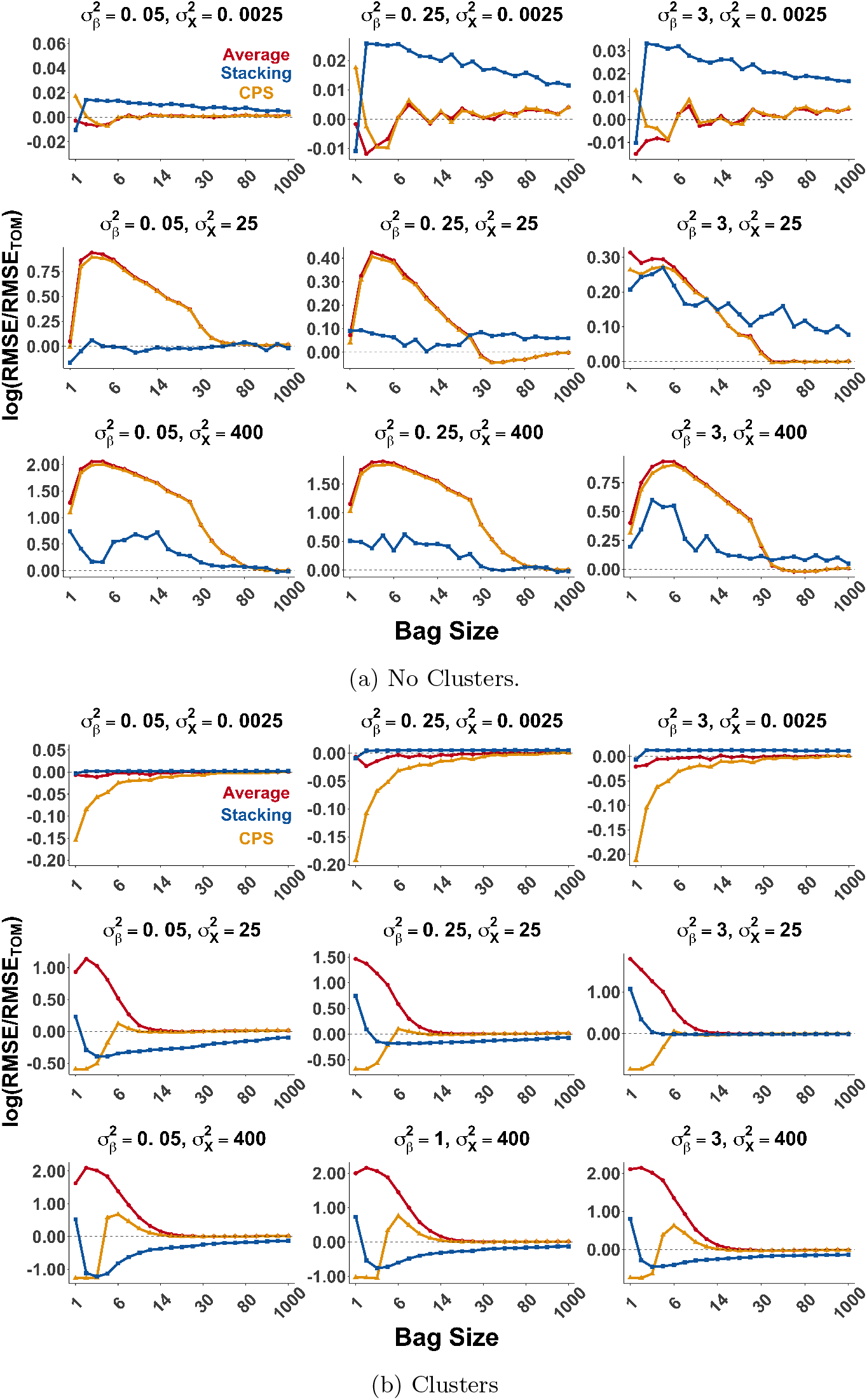
Performance of SSE algorithm on test set as a function of bag size. Relative performance of TOM indicated with black line.

**Fig S.5:**
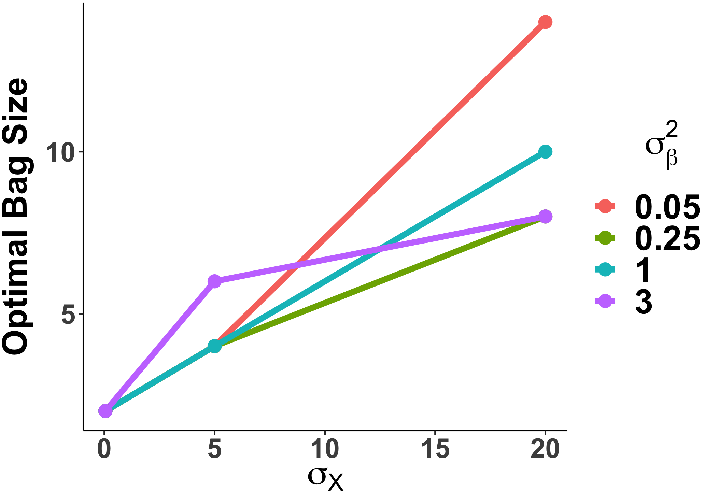
AR algorithm bag size curve: Optimal bag size scales with 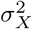. Color indicates 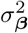. Minor noise was added to points for clarity.

## Appendix C: NEUROSCIENCE DATA

### C.1. Data Description

The data was collected by exposing an electrode in a flow cell (in vitro) to different prepared concentrations (*c*_*j*_) of a neurotransmitter, dopamine, a naturally occurring brain chemical (where the concentration is known to the investigator). Many measurements are taken at a given concentration (these different measurements are referred to as “Replicate” below). While measurements are taken across time, time is ignored in the training set (i.e., each observation is treated as independent). The rows of the dataset then correspond to a measurement (observation) at a given concentration, at a given time point. Furthermore, these data are paired with a vector of labels of known neurotransmitter concentrations ([*DA*]_*i*_). The structure of the data for the *k*^*th*^ electrode is presented in Table 2.

A sample of the average covariates of four of the 15 electrodes is presented in Figure S.6 for both the raw and derivative pre-processing versions. The figure highlights the standardization in covariates that the derivative provides.

#### Similarity Metric

In order to compare the covariates of two studies, we developed a measure that summarizes the covariate profile of a given observation. Our similarity measure was designed based upon the observation that the inflection points in the average CV of each dataset appeared to differ both in magnitude and the in the voltage potential (i.e., the covariate index) at which they occurred. These features are the coordinates of the inflection points of the CVs (Figure S.8b): ***ν*** contains theses coordinates collapsed into a single vector (i.e., ***ν*** ∈ ℝ^8^). Then to compare, the *i*^*th*^ and *j*^*th*^ studies, we calculate the distance between the average ***ν*** of each study via the similarity metric, 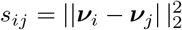 (Figure S.8a). Studies/electrodes differed not only in the average magnitude of the covariates (i.e., the height of the figures), but also the covariates that were concentration-sensitive (i.e., which coefficients, ***β*** were non-zero). This measure was designed to account for this by measuring distance in terms the covariate index and height.

**Fig S.6:**
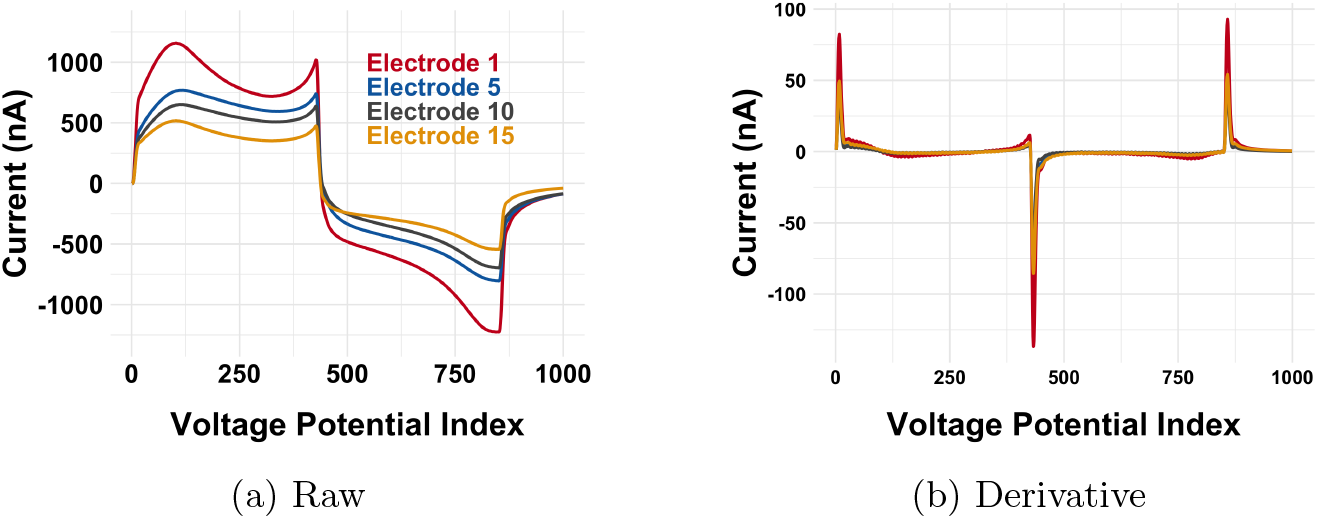
Between-electrode variability: a comparison of average CVs (the covariates) of four different electrodes/studies. The x-axis is the covariate index (i.e., the *j*^*th*^ Voltage Potential is the *j*^*th*^ covariate) and the y-axis is the magnitude of the corresponding covariate. Data are presented here since the 1000 covariates can also be viewed as a single functional covariate. The covariates are all on the same scale and units.

### C.2. Modeling and Methods

We estimated the derivative of the covariates with respect to the voltage potential using the `diff()` function in R, since the measurements were taken at evenly spaced intervals (in time and in voltage potential).

**Fig S.7:**
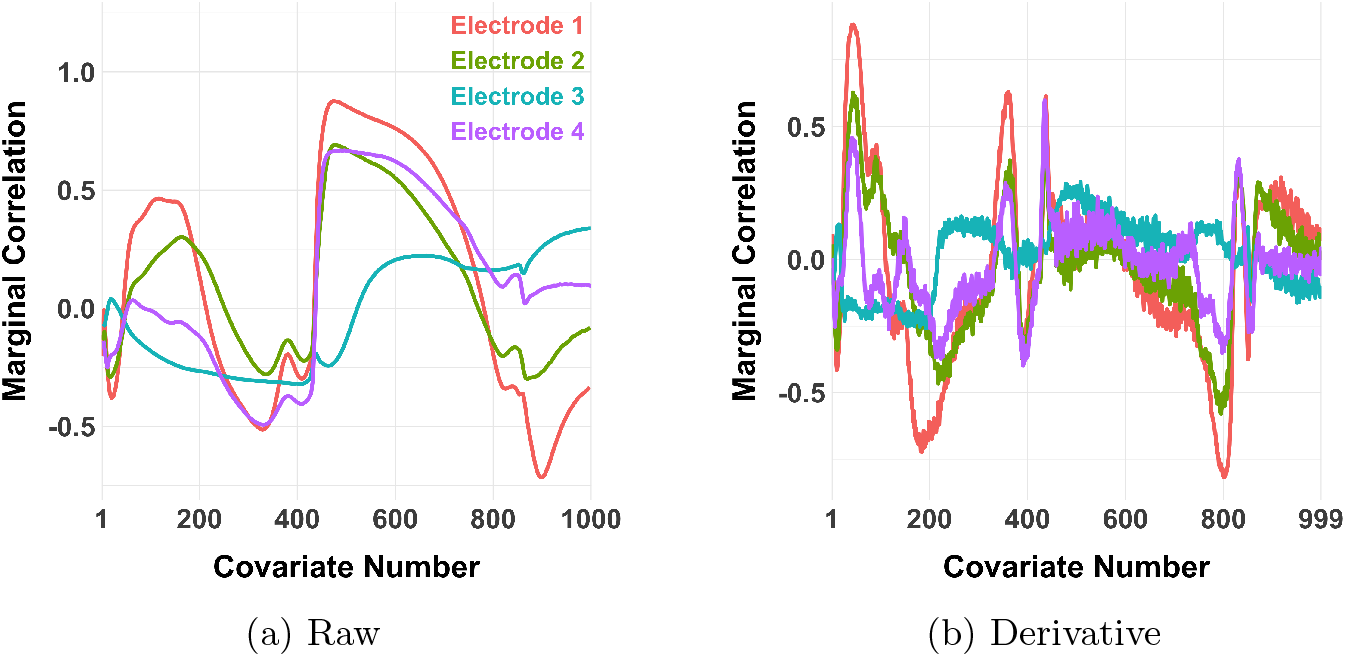
Marginal correlation coefficient estimates 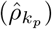 between each covariate and the outcome from four example studies for each covariate.

**Fig S.8:**
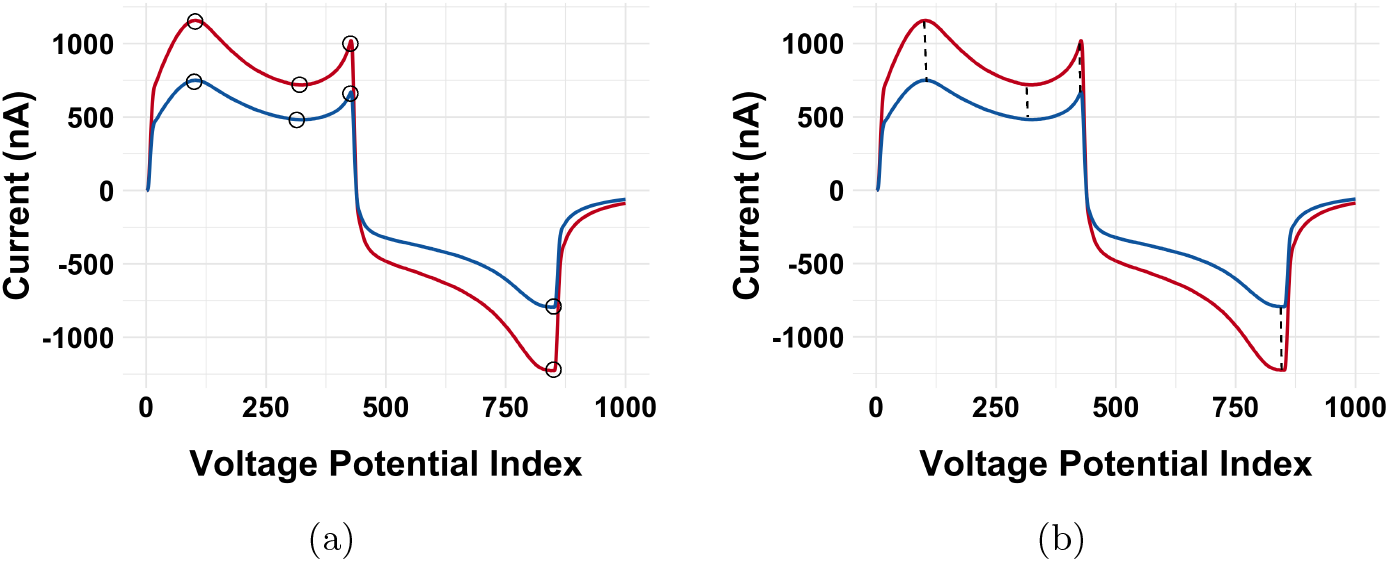
(a) Inflection points of the average of the two studies. (b) Similarity metric: sum of the squared lengths of the diagonal dotted lines.

**Fig S.9:**
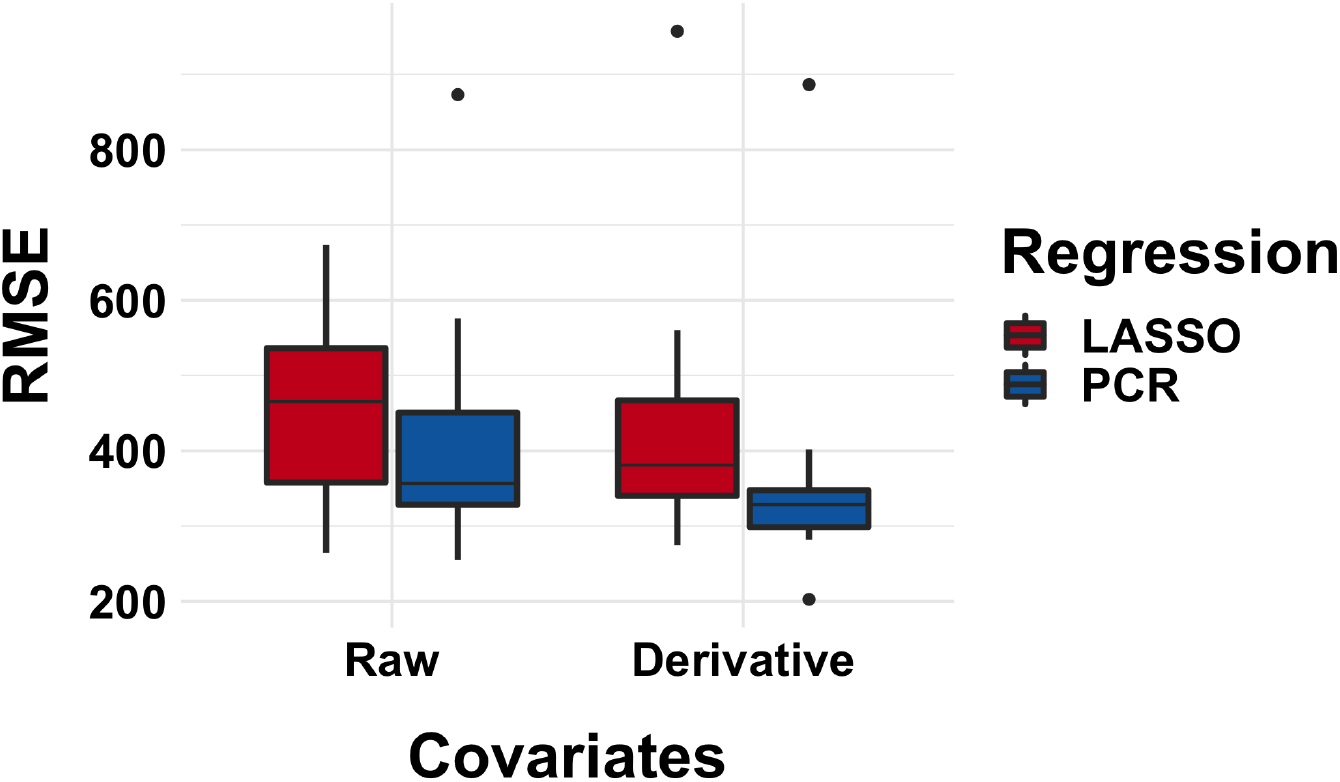
Predictive performance of TOM algorithm for both the raw and derivative data preprocessing. LASSO and PCR hyperparameters both tuned via hold-one-study-out CV.

**Table 2.** Structure of the data for the k^th^ electrode/study

| [DA] | Replicate | $x_1$ | $x_2$ | ⋯ | $x_{1000}$ |
| --- | --- | --- | --- | --- | --- |
| $c_1$ | 1 | $x_{1,1}$ | $x_{1,2}$ | ⋯ | $x_{1,1000}$ |
| $c_1$ | 1 | $x_{2,1}$ | $x_{2,2}$ | ⋯ | $x_{2,1000}$ |
| ⋯ | ⋯ | ⋯ | ⋯ | ⋯ | ⋯ |
| $c_1$ | 700 | ⋯ | ⋯ | ⋯ | ⋯ |
| $c_2$ | 1 | ⋯ | ⋯ | ⋯ | ⋯ |
| $c_2$ | 2 | ⋯ | ⋯ | ⋯ | ⋯ |
| ⋯ | ⋯ | ⋯ | ⋯ | ⋯ | ⋯ |
| $c_2$ | 700 | ⋯ | ⋯ | ⋯ | ⋯ |
| ⋯ | ⋯ | ⋯ | ⋯ | ⋯ | ⋯ |
| ⋯ | ⋯ | ⋯ | ⋯ | ⋯ | ⋯ |
| $c_{30}$ | 700 | $x_{2,1}$ | $x_{2,2}$ | ⋯ | $x_{2,1000}$ |

**Fig S.10:**
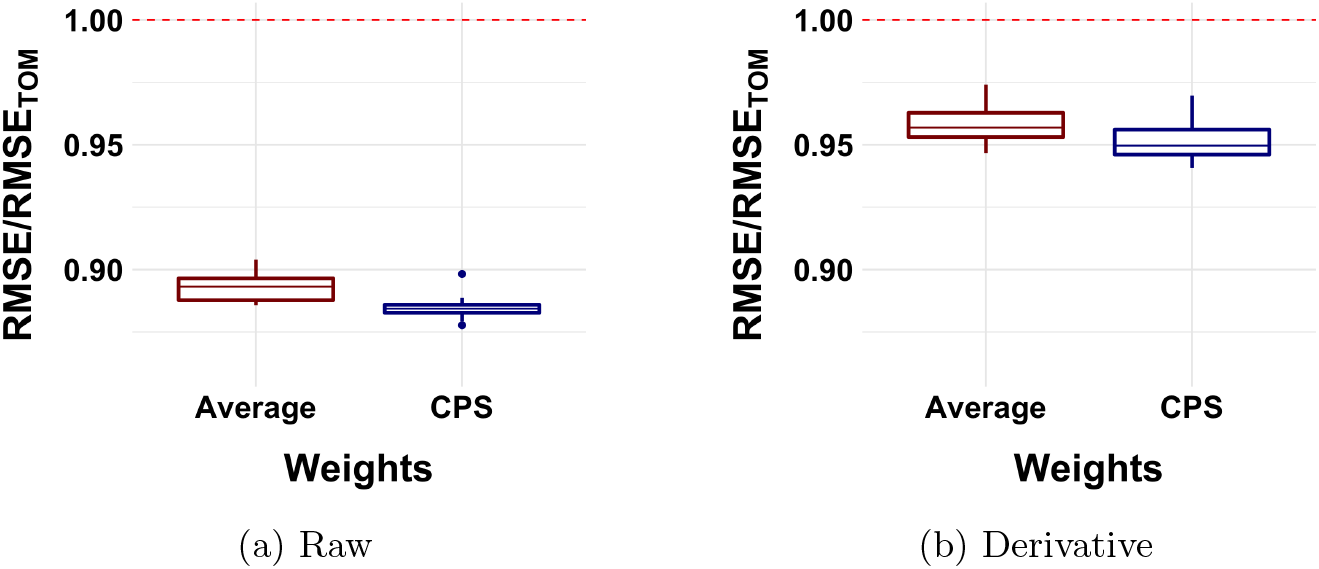
Between-seed variability in average performance (relative to the TOM algorithm). Each point is the mean RMSE/RMSE_*TOM*_ for a single seed, averaged across all held-out-studies.

**Fig S.11:**
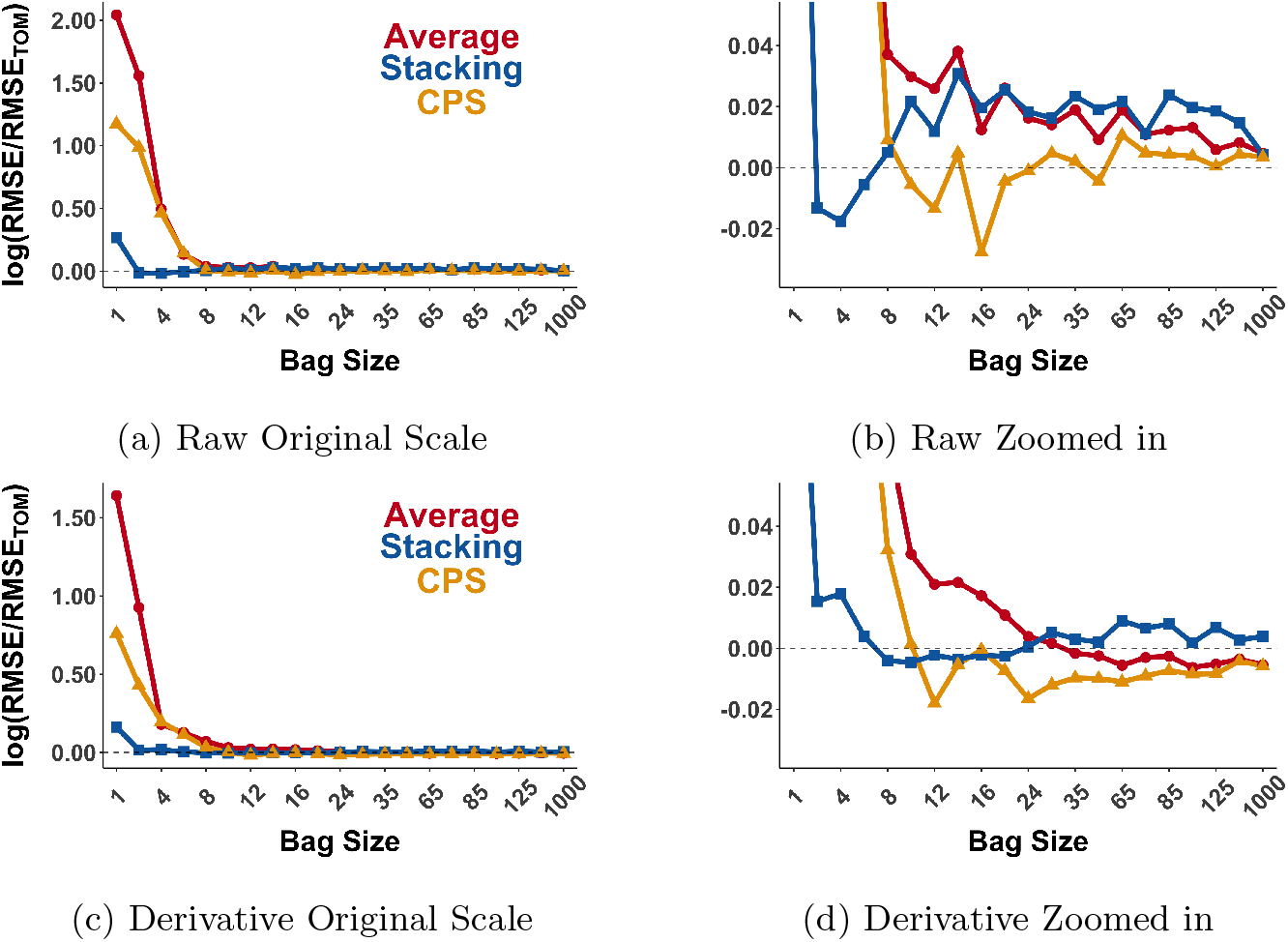
Average Study Strap Ensemble Bag Size Curve: Derivative shifts the optimal bag size to higher values). Vertical lines indicate optimal bag size. RMSEs are standardized to the RMSE of the T OM algorithm fit on the Raw and Derivative respectively.

